# Spatiotemporal limits of optogenetic manipulations in cortical circuits

**DOI:** 10.1101/642215

**Authors:** Nuo Li, Susu Chen, Zengcai V. Guo, Han Chen, Yan Huo, Hidehiko K. Inagaki, Courtney Davis, David Hansel, Caiying Guo, Karel Svoboda

## Abstract

Neuronal inactivation is commonly used to assess the involvement of groups of neurons in specific brain functions. Optogenetic tools allow manipulations of genetically and spatially defined neuronal populations with excellent temporal resolution. However, the targeted neurons are coupled with other neural populations over multiple length scales. As a result, the effects of localized optogenetic manipulations are not limited to the targeted neurons, but produces spatially extended excitation and inhibition with rich dynamics. Here we benchmarked several optogenetic silencers in transgenic mice and with viral gene transduction, with the goal to inactivate excitatory neurons in small regions of neocortex. We analyzed the effects of the perturbations *in vivo* using electrophysiology. Channelrhodopsin activation of GABAergic neurons produced more effective photoinhibition of pyramidal neurons than direct photoinhibition using light-gated ion pumps. We made transgenic mice expressing the light-dependent chloride channel GtACR under the control of Cre-recombinase. Activation of GtACR produced the most potent photoinhibition. For all methods, localized photostimuli produced photoinhibition that extended substantially beyond the spread of light in tissue, although different methods had slightly different resolution limits (radius of inactivation, 0.5 mm to 1 mm). The spatial profile of photoinhibition was likely shaped by strong coupling between cortical neurons. Over some range of photostimulation, circuits produced the “paradoxical effect”, where excitation of inhibitory neurons reduced activity in these neurons, together with pyramidal neurons, a signature of inhibition-stabilized neural networks. The offset of optogenetic inactivation was followed by rebound excitation in a light dose-dependent manner, which can be mitigated by slowly varying photostimuli, but at the expense of time resolution. Our data offer guidance for the design of *in vivo* optogenetics experiments and suggest how these experiments can reveal operating principles of neural circuits.

## Introduction

The cerebral cortex consists of dozens of distinct areas (Dong, 2008; Felleman and Van Essen, 1991; Paxinos and Watson, 1997). Each brain region in turn contains multiple cell types (Rudy et al., 2011; Tasic et al., 2016; Zeng and Sanes, 2017). An important goal in neuroscience is to link dynamics in neural circuits to neural computation and behavior. Much of what we know about localization of cortical function comes from loss-of-function studies. Classically, lesions (Lashley, 1931; Mishkin and Ungerleider, 1982; Newsome and Wurtz, 1988), pharmacological inactivation (Guo et al., 2017; Hikosaka and Wurtz, 1985; Krupa et al., 1999), or cooling (Long and Fee, 2008; Ponce et al., 2008) have been used to silence activity in small (> 1 mm^3^) regions of tissue. The development of optogenetics has allowed rapid and reversible silencing of neuronal activity (Deisseroth, 2015). Concurrently, there has been significant progress in creating genetic access to specific neuronal populations (Luo et al., 2018). Various transgenic Cre driver mouse lines target subtypes of cortical neurons (Gerfen et al., 2013; Gong et al., 2007; Harris et al., 2014; Taniguchi et al., 2011). Crossed with Cre-dependent reporter lines that endogenously express optogenetic effector proteins (Madisen et al., 2015; Madisen et al., 2012; Zhao et al., 2011), and viral delivery methods (Atasoy et al., 2008; Chatterjee et al., 2018; Deverman et al., 2016; Dimidschstein et al., 2016; Luo et al., 2018; Tervo et al., 2016; Wickersham et al., 2007), these technologies have advanced the precision with which manipulation experiments can be carried out.

Optogenetic loss-of-function experiments rely on two schemes (Wiegert et al., 2017). ’*Direct photoinhibition*’ involves light-gated Cl-/H+ pumps that hyperpolarize neurons (Brown et al., 2018; Chow et al., 2010; Chuong et al., 2014; Han and Boyden, 2007; Zhang et al., 2007) and light-gated Cl- channels that produce hyperpolarization and shunting inhibition (Berndt et al., 2014; Govorunova et al., 2015; Wietek et al., 2014). Direct photoinhibition silences genetically-defined neural populations. A second scheme is ‘*ChR-assisted photoinhibition’*, which relies on activation of excitatory channelrhodopsins (ChR) expressed in GABAergic neurons. Photostimulation of the GABAergic neurons potently inhibits local pyramidal neurons and thereby removes excitatory output from the photoinhibited brain region (Cardin et al., 2009; Guo et al., 2014b; Olsen et al., 2012). ChR-assisted photoinhibition has been applied in transgenic mice (such as VGAT-ChR2-EYFP) that express ChR2 in GABAergic neurons (Zhao et al., 2011); this does not require complex crosses and Cre -mediated expression can be used for other genetic labels. Alternatively, the experimenter can choose to express specific variants of ChR (e.g. ChR2 vs. red-shifted ChR (Hooks et al., 2015; Klapoetke et al., 2014; Lin et al., 2013)) in specific GABAergic neurons (e.g. PV vs. SOM neurons) using interneuron-specific Cre lines.

Both direct photoinhibition and ChR-assisted photoinhibition have been widely used in mice to reveal the involvement of brain areas and neuronal populations in specific phases of behavior (Goard et al., 2016; Guo et al., 2015; Guo et al., 2014b; Hanks et al., 2015; Kwon et al., 2016; Li et al., 2015; Li et al., 2016; Mathis et al., 2017; Morandell and Huber, 2017; Resulaj et al., 2018; Sachidhanandam et al., 2013). However, neural circuits have local and long-range connections (Harris and Shepherd, 2015; Hooks et al., 2011; Hooks et al., 2013; Kato et al., 2017; Lefort et al., 2009; Mao et al., 2011; Ozeki et al., 2009; Xue et al., 2014). Strong and nonlinear coupling between neurons, in addition to the photostimulus itself, can affect the spatial and temporal patterns in changes of activity. As a result unexpected effects of optogenetic manipulations are common. For example, activating and inhibiting SOM+ and PV+ neurons can produce complex and asymmetric effects on excitatory neuron stimulus selectivity in visual cortex (Adesnik et al., 2012; Atallah et al., 2012; Lee et al., 2012; Wilson et al., 2012) and auditory cortex (Phillips and Hasenstaub, 2016; Seybold et al., 2015). The effect depends on spontaneous activity, nonlinearity (Phillips and Hasenstaub, 2016), connectivity of the neural circuits (Seybold et al., 2015), or strength and duration of the light manipulation (Atallah et al., 2014; Guo et al., 2014b; Lee et al., 2014). Theoretical studies of networks of excitatory and inhibitory neurons predict counterintuitive responses of neural circuits to perturbations. For example, excitation of cortical interneurons can cause a decrease in activity of the same interneurons (’paradoxical effect’), which arises from the interplay between recurrent excitation and inhibition (Litwin-Kumar et al., 2016; Pehlevan and Sompolinsky, 2014; Rubin et al., 2015; Sadeh et al., 2017; Tsodyks et al., 1997). Optogenetic manipulations of a local region can also impact activity of downstream regions in complex ways. Activity in a brain region can be robust to perturbation of a strongly connected brain region (Li et al., 2016), yet perturbation of other connected brain regions can have a dramatic effect (Guo et al., 2017). Perturbation experiments using behavior as readout must be interpreted in terms of measured changes in neuronal activity. The direct and indirect effects of circuit manipulations on neuronal activity are rarely measured, particularly *in vivo*.

We characterized several optogenetic methods to locally silence somatosensory and motor cortex, either using direct photoinhibition or ChR-assisted photoinhibition. In addition we measured the wavelength-dependent spread of light and its effects on the spatial spread of inactivation. Many light-gated opsins (such as ChR2) are excited by blue light. But blue light is highly scattered and absorbed by blood (Chow et al., 2010; Stujenske et al., 2015; Wiegert et al., 2017; Yizhar et al., 2011; Yona et al., 2016). Red light is less subject to hemoglobin absorption (Svoboda and Block, 1994), and thus can propagate further and produces less local heating (Liu et al., 2015; Stujenske et al., 2015; Wiegert et al., 2017). Red-shifted opsins (e.g. ReaChR (Lin et al., 2013), Chrimson (Klapoetke et al., 2014), Jaws (Chuong et al., 2014)) could enable noninvasive manipulations of deep brain regions. However, direct measurements of light propagation in the intact brain is scarce (Guo et al., 2014b; Ranganathan et al., 2018; Yizhar et al., 2011; Yona et al., 2016), and measurements of the distribution of light intensity are missing. We directly measured light intensity in cortex at two commonly used wavelengths (blue and orange, 473 and 594nm) using a photobleaching assay (Guo et al., 2014b).

We found that the resolution of optogenetic silencing is limited, with silencing spreading beyond the edge of the photostimulus. The length scale of photoinhibition is caused by strong coupling within cortical circuits: loss of activity in a focal region withdraws input to other layers and surrounding regions, resulting in loss of activity in both excitatory and inhibitory populations in the surround, as predicted by network models (Litwin-Kumar et al., 2016; Rubin et al., 2015; Sadeh et al., 2017; Tsodyks et al., 1997). Offset of the photostimulus was typically followed by rebound excitation in a light dose-and duration-dependent manner, which limits the temporal resolution of photoinhibition. Our data outline spatial and temporal constraints of optogenetics manipulations *in vivo* and provide guidance to the design of loss-of-function experiments.

## Results

### Optogenetic inactivation

We examined eight different optogenetic methods to inactivate cortical activity in awake mice (Table 1, Fig. 1A).

**Table 1.** A list of photoinhibition methods tested in this study.

| Methods | Mouse (JAX #) | Reagents | Wavelength | Brain region |
| --- | --- | --- | --- | --- |
| ***ChR-assisted photoinhibition*** |  |  |  |  |
| ChR2 in all GABAergic neurons | VGAT-ChR2-EYFP (014548) |  | 473 nm | Somatosensory cortex, ALM |
| ChR2 in PV neurons | PV-IRES-Cre (008069) x Ai32 (012569) |  | 473 nm | Somatosensory cortex |
| ChR2 in SOM neurons | SOM-IRES-Cre (013044) x Ai32 (012569) |  | 473 nm | Somatosensory cortex |
| ReaChR in PV neurons | PV-IRES-Cre (008069) x R26-LSL-CAG-LSL-ReaChR-mCit (026294) |  | 594 nm | ALM |
| ChR2 virally delivered to local PV neurons | PV-IRES-Cre (008069) | AAV2/1-CAG-FLEX-ChR2-tdTomato-WPRE (UPenn Viral Core, AV-1-ALL864) | 473 nm | Somatosensory cortex |
| ***Direct photoinhibition*** |  |  |  |  |
| Arch in excitatory neurons | Emx1-Cre (005628) x Ai35D (012735) |  | 594 nm | Somatosensory cortex |
| Jaws in excitatory neurons | Emx1-Cre (005628) x Ai79D (023529) x Camk2-tTA (003010) |  | 594 nm | Somatosensory cortex |
| GtACR1 (somatic targeting) (Mahn et al., 2018) in excitatory neurons | Emx1-Cre (005628) x R26-CAG-LNL-GtACR1-ts-FRed-Kv2.1 (033089) |  | 473 nm, 635 nm | Somatosensory cortex, primary motor cortex, ALM |

**Figure 1.**
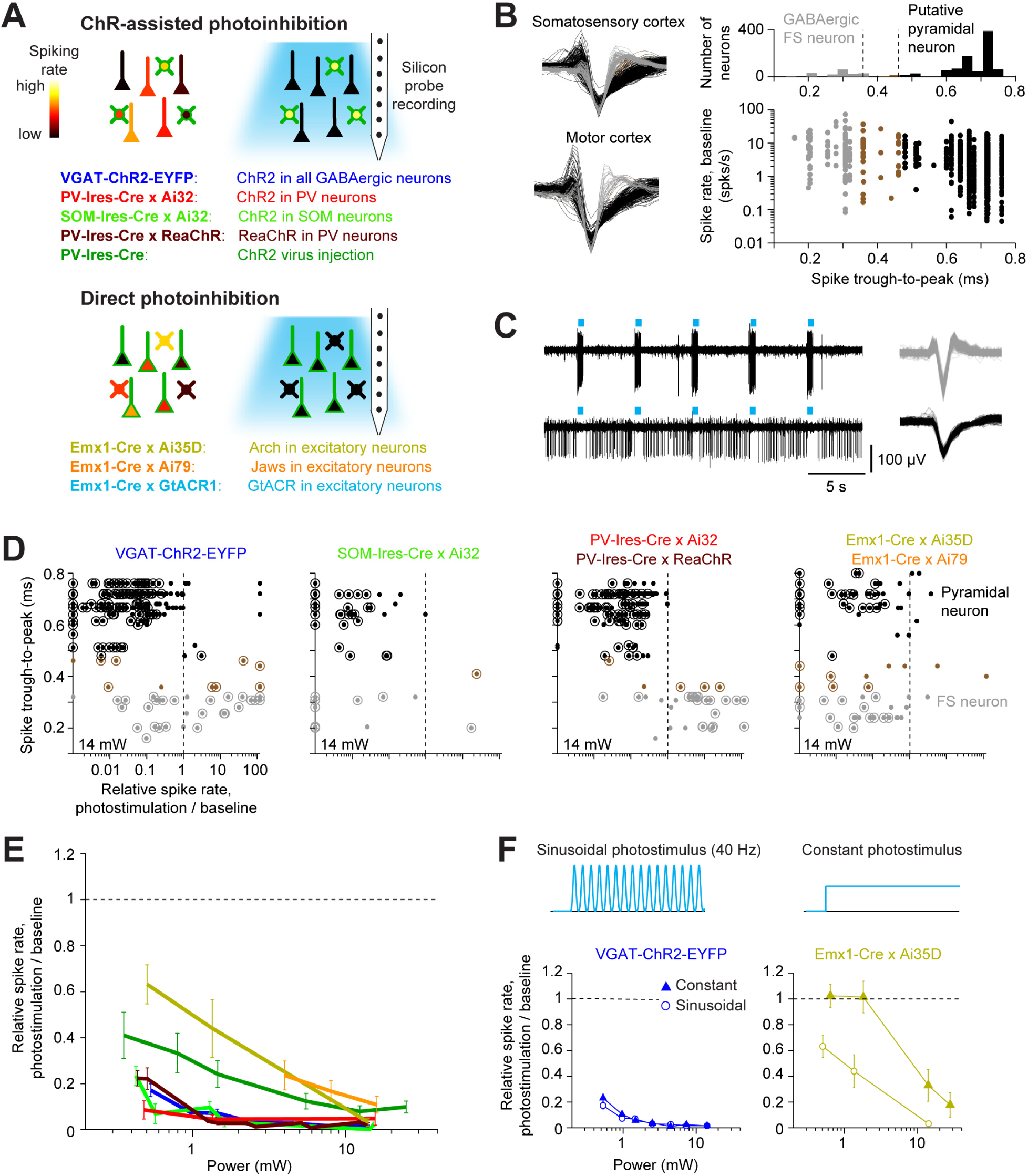
Optogenetic inactivation and cell-type specific recording. (A) Different inactivation methods. ChR2-assisted photoinhibition was induced by photostimulating ChR2 in various GABAergic interneurons. Direct photoinhibition was achieved by photostimulating inhibitory opsins in pyramidal neurons. (B) Silicon probe recordings. Cell-type classification based on spike waveform. Left, spike waveforms for putative FS neurons (gray) and pyramidal neurons (black) in two different cortical areas. Right, histogram of spike durations. Neurons that could not be classified based on spike width were excluded from analysis (brown; see Materials and Methods). (C) Silicon probe recordings during photostimulation in a VGAT-ChR2-EYFP mouse. Top, a GABAergic fast spiking (FS) neuron. Bottom, a putative pyramidal neuron. Right, corresponding spike waveforms. (D) Effects of photostimulation. Responses were normalized to baseline (dashed line). Dots correspond to individual neurons. Neurons with significant firing rate changes relative to baseline (*p* < 0.05, two-tailed *t*-test) are highlighted by circles. Same color scheme as (B). (E) Spike rate as a function of laser power (< 0.4 mm from laser center, all cortical depths). Pyramidal neurons only. Spike rates were normalized to baseline (dashed line, see Experimental Procedures). Mean ± s.e.m. across neurons, bootstrap. Same color scheme as (A). VGAT-ChR2-EYFP, n=153; PV-IRES-Cre x Ai32, n=16; SOM-IRES-Cre x Ai32, n=65; PV-IRES-Cre x ReaChR, n=211; PV-IRES-Cre, ChR2 virus injection, n=78; Emx1-Cre x Ai35D, n=26; Emx1-Cre x Camk2a-tTA x Ai79, n=176. (F) Top, photostimuli. Standard photostimulus (40 Hz Sinusoid) and constant photostimulus. Bottom, spike rate as a function of laser power (<0.4 mm from laser center, all cortical depths) for ChR-assisted photoinhibition (left) and direct photoinhibition mediated by Arch (right).

For ChR-assisted photoinhibition we photostimulated excitatory opsins in GABAergic interneurons to drive inhibition in nearby pyramidal neurons. We used transgenic mice that expressed ChR2 in all GABAergic neurons (VGAT-ChR2-EYFP), or in parvalbumin-positive (PV) interneurons (PV-IRES-Cre X Ai32), or in somatostatin-positive (SOM) interneurons (SOM-IRES-Cre X Ai32). In addition, we photostimulated a red-shifted channelrhodopsin (ReaChR) in PV neurons (PV-IRES-Cre X ReaChR) (Hooks et al., 2015; Lin et al., 2013). We also induced photoinhibition with a Cre-dependent ChR2 virus in PV-IRES-Cre mice. For direct photoinhibition we photostimulated inhibitory opsins in pyramidal neurons, including the ion pumps Archaerhodopsin (Arch, Emx1- Cre X Ai35D) and Jaws (Emx1-Cre X Camk2a-tTA X Ai79) (Chow et al., 2010; Chuong et al., 2014; Madisen et al., 2015).

We measured neural activity in the vicinity of the photostimulus using silicon probe recordings. Measurements were performed in somatosensory cortex and motor cortex; these two cortical regions are examples of sensory and frontal cortex, which differ in laminar connectivity (Hooks et al., 2011) and in their connections with thalamus (Hooks et al., 2013) (Materials and Methods). In both brain areas the distribution of single-unit spike width was bimodal (Fig. 1B). Units with narrow spikes were fast spiking (FS) neurons and likely expressed parvalbumin (Cardin et al., 2009; Guo et al., 2014b; Olsen et al., 2012; Resulaj et al., 2018). Neurons with wide spikes were likely mostly pyramidal neurons, and we refer to this population as putative pyramidal neurons.

These classifications were consistent with spike rate changes observed during photostimulation (Fig. 1C). In mice expressing excitatory opsins in FS GABAergic neurons (VGAT-ChR2-EYFP, PV-IRES-Cre X Ai32, and PV-IRES-Cre X ReaChR mice), FS neurons were activated by photostimulation, whereas putative pyramidal neurons were inhibited (Fig. 1D). In VGAT-ChR2-EYFP mice a subset of FS neurons were inhibited rather than excited, likely caused by activation of other GABAergic neurons that were photostimulated in these mice. In SOM-IRES-Cre X Ai32 mice, both FS neurons and putative pyramidal neurons were inhibited, consistent with SOM neurons inhibiting both PV and pyramidal neurons (Lee et al., 2013; Pfeffer et al., 2013). SOM neurons were rare in extracellular recordings, with only two neurons showing clear photostimulus-induced increases in spike rate. Putative SOM neurons could not be unambiguously separated from either FS neurons or putative pyramidal neurons based on their spike waveforms (Fig. 1D). In Emx1-Cre X Ai35D and Emx1-Cre X Camk2a-tTA X Ai79 mice, optogenetic hyperpolarization of pyramidal neurons reduced activity of putative pyramidal neurons and FS neurons. The inhibition of FS neurons was likely caused by withdrawal of excitatory input from pyramidal neurons.

We measured the light sensitivity of different optogenetic inactivation methods by normalizing the spike rates of putative pyramidal neurons to their baseline spike rates (Materials and Methods). Near the center of the photostimulus (< 0.4 mm from laser center), ChR-assisted photoinhibition in transgenic mice produced strong inactivation, with > 80% of reduction in spike rates for putative pyramidal neurons over a wide range of laser powers (0.5-10mW, Fig 1E). ChR2 virus mediated photoinhibition in PV-IRES-Cre mice produced slightly weaker silencing. Arch and Jaws-mediated photoinhibition required roughly 10-fold higher laser power to achieve similar silencing compared to photoinhibition (Fig 1E). For Arch-mediated photoinhibition, temporally-modulated photostimuli were more effective in silencing activity than constant photostimuli at the same average power (Fig 1F).

We also photoactivated the *Guillardia theta* anion channel rhodopsin 1 (GtACR1) to hyperpolarize and shunt the membranes of pyramidal neurons (Govorunova et al., 2015). We generated a Cre reporter mouse, expressing a soma localized GtACR1 (Mahn et al., 2018) driven by the CAG promoter targeted to the Rosa26 locus (Hooks et al., 2015; Madisen et al., 2010; Muzumdar et al., 2007) (Fig 2A) (R26-CAG-LNL-GtACR1-ts-FRed-Kv2.1, JAX #033089). We expressed soma localized GtACR1 in cortical excitatory neurons by crossing the reporter mouse to Emx1-Cre. GtACR1 expression was targeted to the soma, but weak expression was still seen in axons (Fig 2A, 2B). Under blue light (473 nm) illumination, GtACR1 produced the most potent photoinhibition near the laser center, with > 80% of the spikes silenced at 0.1 mW (Fig 2C). Previous studies have shown that GtACR1 and other Cl-channels induce spiking in axons, caused by a positively shifted chloride reversal potential in the axon (Mahn et al., 2018; Messier et al., 2018). At moderate laser power (0.2 mW), photostimulation of soma localized GtACR1 induced little axonal excitation (Mahn et al., 2018; Messier et al., 2018) (Fig 2D) although at higher laser powers axonal excitation was still apparent. GtACR1 mediated photoinhibition could also be induced using red light (Fig 2C, 635 nm) (albeit at higher light intensities), which could facilitate photoinhibition of deep brain regions. Expression of GtACR1 appears to be by far the most sensitive method for inactivation, although attention has to be paid to avoid axonal excitation.

**Figure 2.**
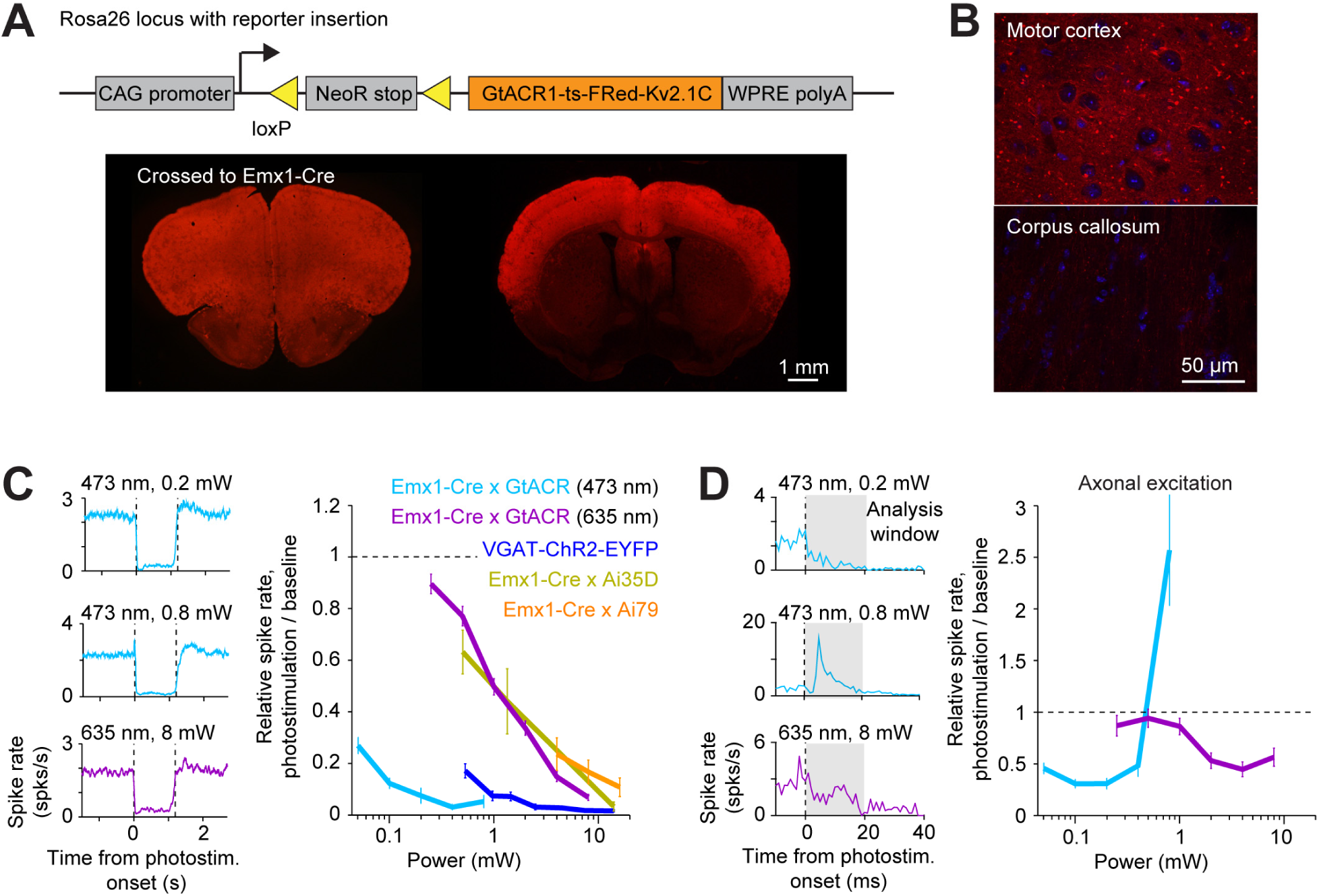
Direct photoinhibition with a GtACR1 reporter mouse. (A) *Top*, generation of the Cre-dependent GtACR reporter line. Construct includes loxP sites, and GtACR1-ts-FRed-Kv2.1C with WPRE, driven by CAG promoter targeted to the Rosa26 locus. *Bottom*, cross to Emx1-Cre mice. GtACR1 is expressed in cortical excitatory neurons. (B) Confocal images showing dense expression of GtACR1 in cortex and low levels of expression in the corpus callosum and thalamus, implying low trafficking to axons. (C) *Left*, mean peristimulus time histogram (PSTH, 50 ms bin) for pyramidal neurons with blue and red photostimuli. All neurons < 0.4 mm from the laser center were pooled. Blue light photostimulation, n=335 neurons from ALM; Red light photostimulation, n=285 neurons from ALM. *Right*, normalized spike rate as a function of laser power. Mean ± s.e.m. across neurons, bootstrap. VGAT-ChR2-EYFP, Emx1-Cre x Ai35D, Emx1-Cre x Camk2a-tTA x Ai79, data from Figure 1E replotted here for reference. (D) *Left*, PSTH (1 ms bin) at the onset of the photostimulation. Axonal excitation was induced only at 0.8 nW laser power. Right, normalized spike rate during the first 20 ms of photostimulation onset.

## Spatial profile of light intensity

To characterize the spread of inactivation we first measured the spatial profile of the photostimulus, that is, the light intensity in the tissue. At the surface of the brain the photostimulus was a laser beam with a Gaussian profile (diameter at 4σ, 400 µm), that was focused onto the cortical surface through a clear skull implant (Materials and Methods). In the brain, light is scattered and absorbed, primarily by blood. As a substitute for light intensity we measured the three-dimensional profile of photobleaching of fluorescent proteins in transgenic mice. Because neurons are large (100’s of micrometers) and fluorescent proteins diffuse rapidly in the cytoplasm (Swaminathan et al., 1997), we used fluorescent proteins targeted to neuronal nuclei. We measured the spatial distribution of fluorescence after prolonged light exposure, which caused pronounced photobleaching at the center of the photostimulus. For blue light (473nm), we used transgenic mice expressing GFP in the nuclei of cortical excitatory neurons (Rosa-LSL-H2B-GFP crossed to Emx1-Cre) (He et al., 2012). For orange light (594 nm), we used transgenic mice expressing mCherry (Rosa-LSL-H2B-mCherry crossed to Emx1-Cre) (Peron et al., 2015). Blue light induced photobleaching in a confined region near the photostimulus (Fig 3A). Photobleaching increased with the light dose (Fig 3B). We measured photobleaching by averaging the fluorescence change (ΔF/ F_0_) within a small region near the photostimulus center relative to the baseline fluorescence in regions far away from the laser center (F_0_, Fig 3C). Using an empirical relationship between bleaching and the light dose (light intensity x time) we inferred the spatial profile of light intensity in tissue (Fig 3D, Materials and Methods). Light intensity attenuated to less than 20% at 500 µm below the surface of the brain (Fig 3D; depth at half max, 300 µm). Laterally, light intensity dropped off rapidly (radius at half max, 135 µm) (Fig 3D). These data show that blue light is highly confined in cortical tissue, consistent with previous simulation results (Stujenske et al., 2015).

**Figure 3.**
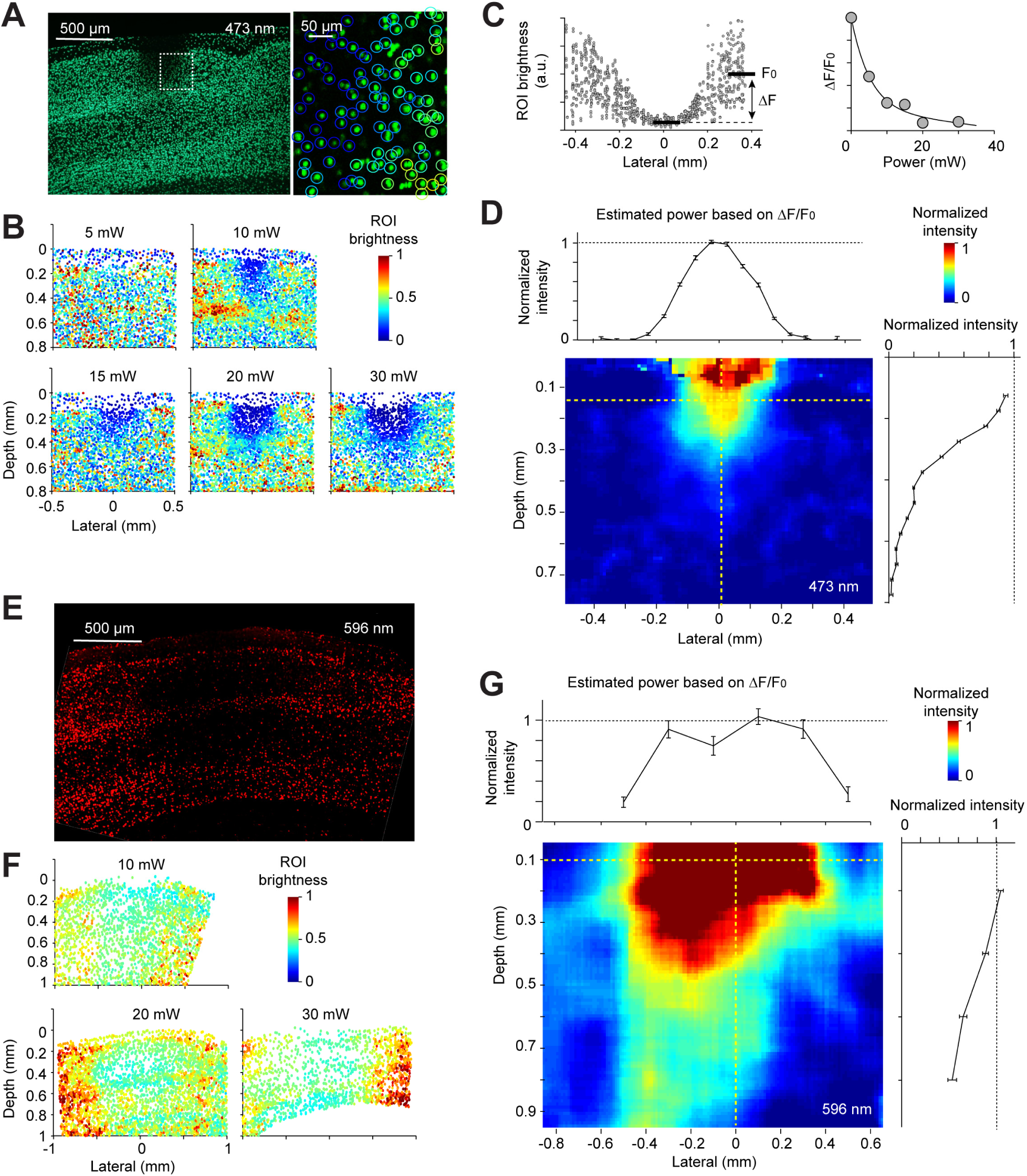
Spatial profile of light in the cortex. (A) Photobleaching was induced in a transgenic mouse line expressing GFP in the nuclei of excitatory neurons (Rosa-LSL-H2B-GFP crossed to Emx1-Cre). *Left*, coronal section showing an example photobleaching site. Light dose, 15 mW, 600 seconds. The photostimulus was a laser beam with a Gaussian profile (4σ = 0.4 mm) at the brain surface. Right, a confocal image showing a region highlighted by the dash line. GFP intensity was measured in regions of interest (ROI) around individual nuclei (circles). Color indicates GFP intensity. (B) Photobleaching assay measuring the spread of blue light in the neocortex. Photobleaching was induced with different light doses (600 s, powers as indicated; Materials and Methods). Dots, individual nuclei. Color indicates GFP intensity. (C) *Left*, nuclear intensity as a function of lateral distance from the laser center (data from 20mW). F_0_, baseline ROI intensity, computed by averaging all ROIs far away from the laser center. ΔF, ROI intensity change caused by photobleaching, computed by averaging all ROIs near the laser center and subtracting F0 (Materials and Methods). *Right*, photobleaching (ΔF/ F0) at various laser powers. (D) The estimated spatial profile of blue light in tissue. Changes in fluorescence (ΔF/ F0) across different laser powers in (C) allowed conversions from GFP intensity to laser power (Material and Methods). Light intensity is shown as a function of cortical depth (left) and lateral distance (right) from the laser center. (E) - (G) Same as (A) - (D), but for orange light (594nm) measured with photobleaching of mCherry (Material and Methods).

In contrast to blue light, orange light propagated much further (depth at half max, 846 µm), penetrating all layers of cortex and illuminating a cubic millimeter of volume (Fig 3E-G), 28 times larger than for blue light. Because orange light (594 nm) is still absorbed by hemoglobin (Svoboda and Block, 1994), longer wavelength light (e.g. 630 nm) will penetrate substantially deeper into tissue than orange light.

### Spatial profile of optogenetic inactivation

We measured photoinhibition across cortical layers when photostimulating GABAergic neurons using blue or orange light (Fig 4A-D). Recordings were made from the barrel cortex near the center of the photostimulus. Photostimulation of ChR2 in GABAergic neurons produced nearly uniform inhibition of pyramidal neurons across cortical layers (Fig 4B, VGAT-ChR2-EYFP, SOM-IRES-Cre X Ai32, and PV-IRES-Cre X Ai32), despite limited penetration of blue light in tissue (Fig 3). Photoinhibition removed the majority of spikes (> 80%) in a cortical column across a wide range of laser powers (1.5 – 14 mW). A notable exception was seen at low laser powers (0.5mW) in SOM-IRES-Cre X Ai32 mice. Selective excitation of SOM neurons produced a disinhibition of excitatory neurons around layer 4 (Fig 4B, SOM-IRES-Cre X Ai32). This disinhibition in layer 4 was likely mediated by SOM neurons that inhibit PV neurons. Activation of SOM neurons therefore removed a major source of inhibition onto excitatory neurons (Xu et al., 2013). Photostimulation of GABAergic neurons using orange light also induced uniform inhibition across cortical layers (Fig 4C-D, PV-IRES-Cre X ReaChR). Despite different propagation of blue and orange light in tissue (Fig. 3), ChR-assisted photoinhibition was remarkably similar for both illumination wavelengths. At moderate laser power (1.5 mW), blue light excited FS neurons mainly in superficial layers (Fig 4E- F). Loss of activity in superficial layers then silenced the deep layers. At moderate laser power, direct photoinhibition of excitatory neurons using GtACR1 or Jaws also inhibited activity across all layers (Fig 4G-L). Remarkably, the profile of silencing mediated by GtACR1 was also similar for blue and red light (Fig 4G-J).

**Figure 4.**
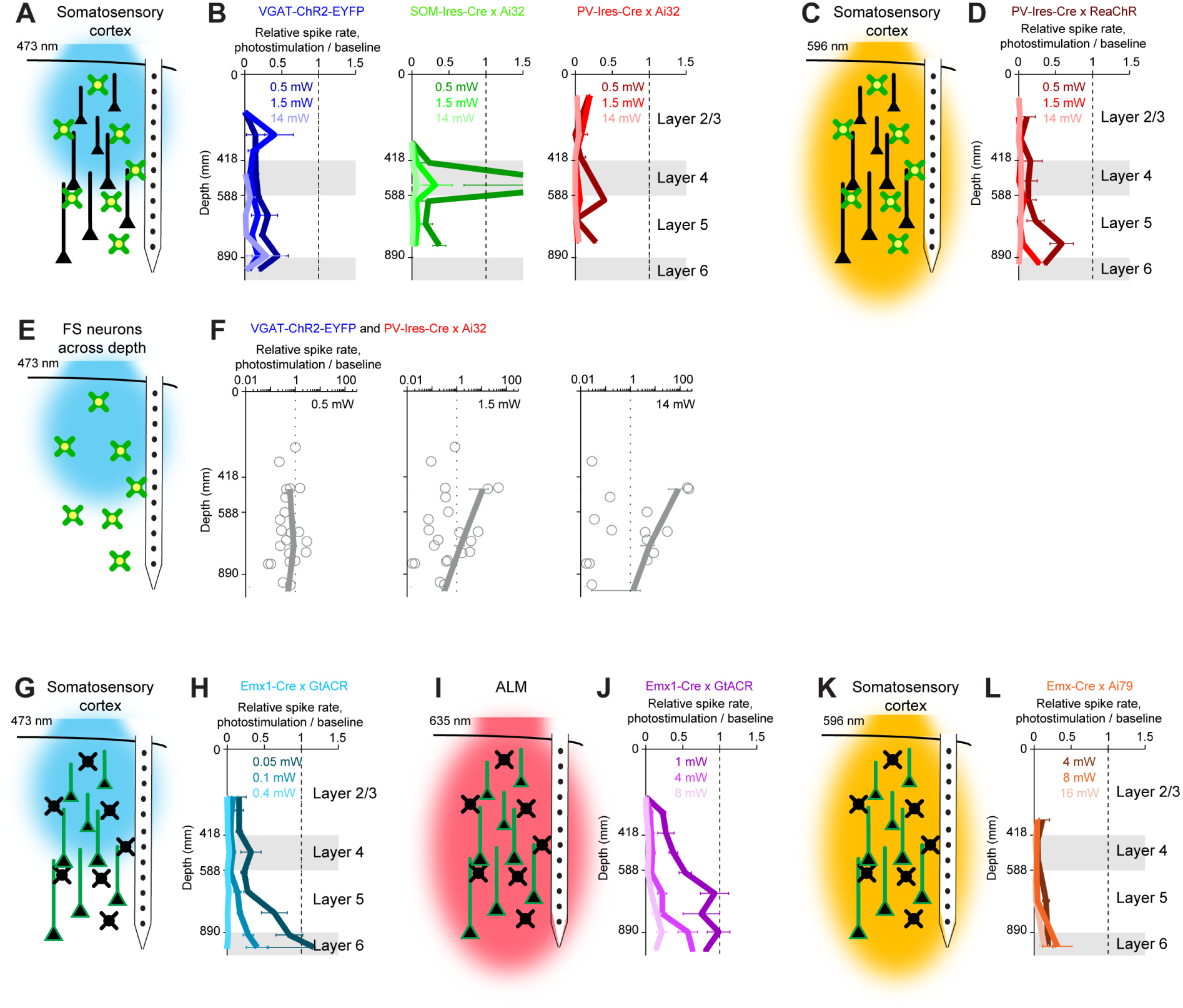
Photoinhibition across cortical layers. (A) ChR-assisted photoinhibition using blue light. (B) Normalized spike rate across cortical depth (<0.2 mm from laser center) for three laser powers (0.5, 1.5, 14 mW). Data from barrel cortex. Pyramidal neurons only. VGAT-ChR2-EYFP, photostimulating all GABAergic neurons (data from 170 pyramidal neurons); SOM-IRES-Cre x Ai32, photostimulating SOM neurons (n=33); PV-IRES-Cre x Ai32, photostimulating PV neurons (n=61). Error bars show s.e.m. over neurons. (C) Schematics, ChR-assisted photoinhibition using orange light. (D) Same as (B), but for red-shifted ChR in PV neurons (PV-IRES-Cre x R26-CAG-LSL-ReaChR-mCitrine, n=95). (E) - (F) Normalized spike rate for FS neurons across cortical depth (<1 mm from laser center) for three laser powers (0.5, 1.5, 14 mW). Data from barrel cortex and motor cortex are combined. Data from VGAT-ChR2-EYFP and PV-IRES-Cre x Ai32 mice are combined (n=22). Dots, individual neurons; mean ± s.e.m. over neurons. (G)- (F) Same as (C) - (D), but for blue light photostimulation of light-gated Cl- channel GtACR1 in pyramidal neurons (Emx1-Cre X GtACR, n=198). (H)- (H) Same as (C) - (D), but for red light photostimulation of GtACR1 in pyramidal neurons (n=236). (I) - (J) Same as (C) - (D), but for light-gated Cl^-^ pump Jaws in pyramidal neurons (Emx1-Cre X Camk2a-tTA X Ai79, n=176).

We next characterized the lateral extent of photoinhibition by varying the location of the photostimulus relative to the recording site (Fig 5A). In VGAT-ChR2-EYFP mice, FS neurons near the laser center were activated in a dose-dependent manner (Fig 5B). The lateral spread of FS neuron excitation was dependent on laser power (at 0.5 mW, half-max width, 0.15 mm; at 14 mW, 1.5 mm, Fig 5B). Photoinhibition of pyramidal neurons extended 1 mm beyond the activation profile of FS neurons (Fig 5B). For example, at 0.5 mW, pyramidal neurons were silenced 1 mm away from the laser center where FS neurons were not activated.

**Figure 5.**
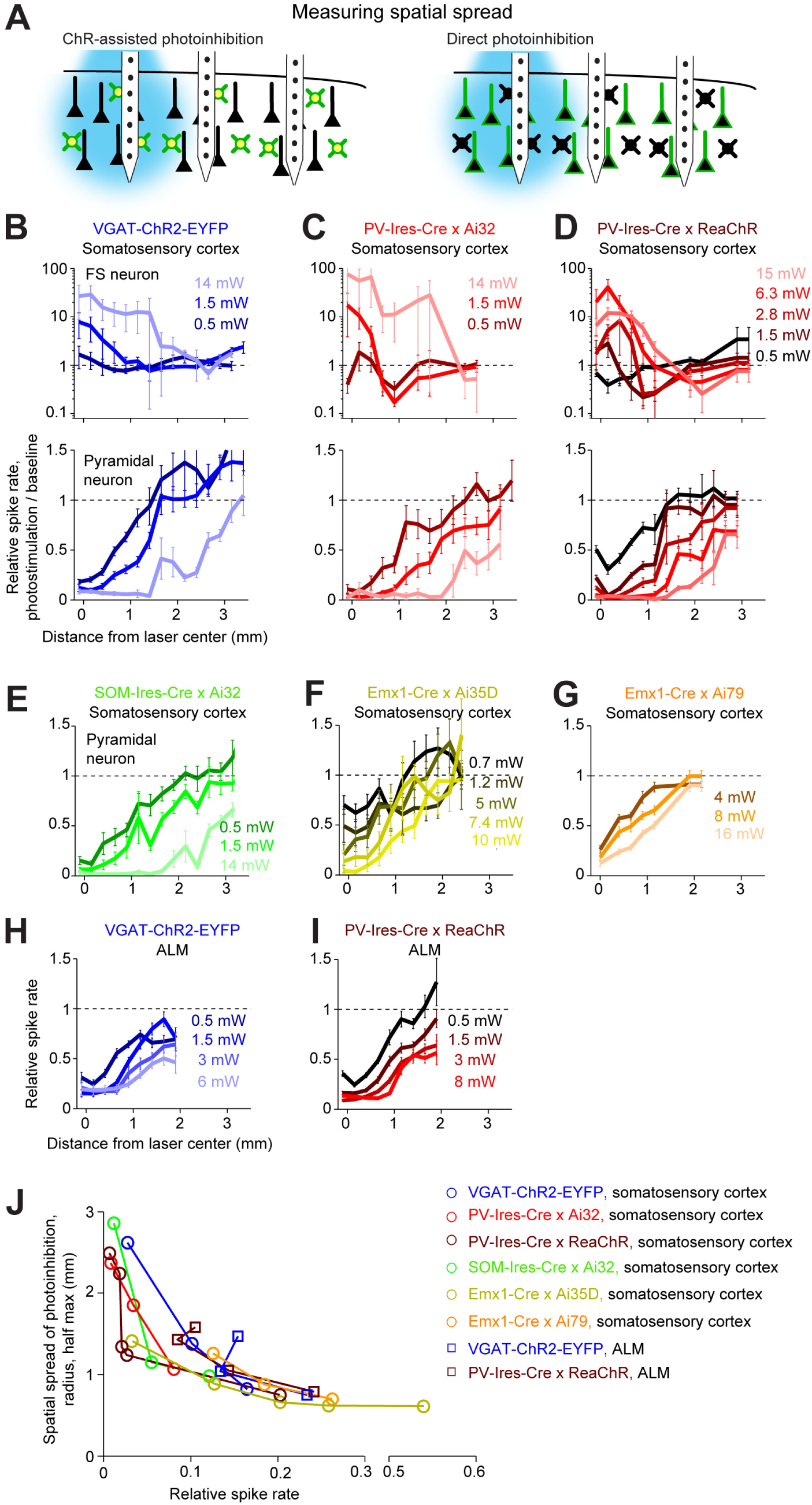
Spatial profile of photoinhibition. (A) Silicon probe recording at different distances from photostimulus. (B) Normalized spike rate versus distance from the photostimulus center for three laser powers (0.5, 1.5, 14 mW). Photostimulation in barrel cortex of VGAT-ChR2-EYFP mice. Top, FS neurons (n=18). Bottom, pyramidal neurons (n=111). Neurons were pooled across cortical depths. Mean ± sem, bootstrap across neurons. (C) Same as (B), photostimulation in PV-IRES-Cre x Ai32 mice (FS neurons, n=5; pyramidal neurons, n=16). (D) Same as (B), photostimulation in PV-IRES-Cre x R26-CAG-LSL-ReaChR-mCitrine using orange light (FS neurons, n=10; pyramidal neurons, n=82). (E) Same as (B), photoinhibition in SOM-IRES-Cre x Ai32 mice. Pyramidal neurons only (n=65). (F) Same as (E), photostimulation in Emx1-Cre x Ai35D with orange light (n=26). (G)Same as (E), photostimulation in Emx1-Cre X Camk2a-tTA X Ai79 with orange light (n=174). (H) Same as (E), photostimulation in anterior lateral motor cortex (ALM) of VGAT-ChR2-EYFP mice (n=96). (I) Same as (E), photostimulation in anterior lateral motor cortex (ALM) of PV-IRES-Cre x R26-CAG-LSL-ReaChR-mCitrine using orange light (n=129) (J) Photoinhibition strength versus spatial spread. Normalized spike rate was averaged across all neurons near laser center (< 0.4 mm, all cortical depths). Spatial spread is the distance at which photoinhibition strength was half of that at the laser center (“radius, half-max”). Each dot represents data from one photostimulation power. Lines connect all dots of one condition.

This spread of photoinhibition extends beyond typical sizes of dendritic and axonal arbors of GABAergic neurons (Jiang et al., 2015) and dendritic arbors of pyramidal neurons (Oberlaender et al., 2012; Oswald et al., 2013; Shepherd et al., 2005; Yamashita et al., 2018). In VGAT-ChR2-EYFP mice, ChR2 was expressed in all GABAergic neurons. We considered the possibility that the spread of photoinhibition was mediated by a specific subtype of GABAergic neuron. Among GABAergic interneurons, PV neurons have the most compact dendritic and axonal arbors (Jiang et al., 2015). We thus tested whether selectively photostimulating PV neurons would produce more spatially-restricted photoinhibition. However, a similarly broad photoinhibition profile was induced in PV-IRES-Cre X Ai32 mice (Fig 5C). We next sought to produce a broader photoinhibition by using orange light to illuminate a larger cortical volume in transgenic mice expressing ReaChR in PV neurons. However, the resulting photoinhibition profile was remarkably similar to those produced by photostimulating ChR2 in PV neurons using blue light (Fig 5D).

We examined other methods of optogenetic inactivation. First, photoinhibition by selective excitation of SOM neurons induced similarly broad photoinhibition (Fig. 5E). Next, direct photoinhibition of pyramidal neurons using light-gated ion pumps (Emx1-Cre X Ai35D mice or Emx1-Cre X Camk2a-tTA X Ai79 mice) induced similarly broad photoinhibition (Fig 5F & 5G). ChR-assisted photoinhibition in a different brain region (anterior lateral motor cortex, ALM) produced similarly broad spatial spread regardless of the method used to drive photoinhibition (Fig 5H-I, blue light photostimulation in VGAT-ChR2-EYFP mice, or orange light photostimulation in PV-IRES-Cre X ReaChR mice). Across all methods, the fractional reduction in spike rate near the laser center approximately predicted the spatial spread of photoinhibition (Fig 5J). These data indicate that, for relatively localized photostimuli, the spatial resolution of photoinhibition laterally and in axially is primarily shaped from interactions between spatially extended activation of GABAergic interneurons with intrinsic properties of cortical circuits.

Higher spatial resolution (i.e. complete removal of spikes from smaller tissue volumes) is desirable for an inactivation method. If the length scale of inactivation arises from intrinsic properties of cortical circuits, it would pose a fundamental constraint on the spatial resolution of neuronal inactivation. Our results suggest that silencing a small area also reduces activity in surrounding brain areas (Fig 5J). Considering the size of the mouse brain (10 mm long) this constraint on resolution is substantial. We looked for conditions that could break the relationship between photoinhibition strength and spatial spread. Transgenic mice express ChR in all GABAergic neurons. At high laser power, photostimulus activates interneurons over a larger area due to light scattering (Fig 3, 4B-D). We next tested the spatial spread of direct photoinhibition induced by GtACR1. In three different cortical regions, GtACR1-mediated photoinhibition also produced broad photoinhibition (Fig 6A), but the spatial spread was slightly lower than that produced by other photoinhibition methods (Fig 6B): with 90% spike rate reduction near the laser center, the radius of silencing was 0.8 mm (;>1 mm for other methods, Fig 8).

**Figure 6.**
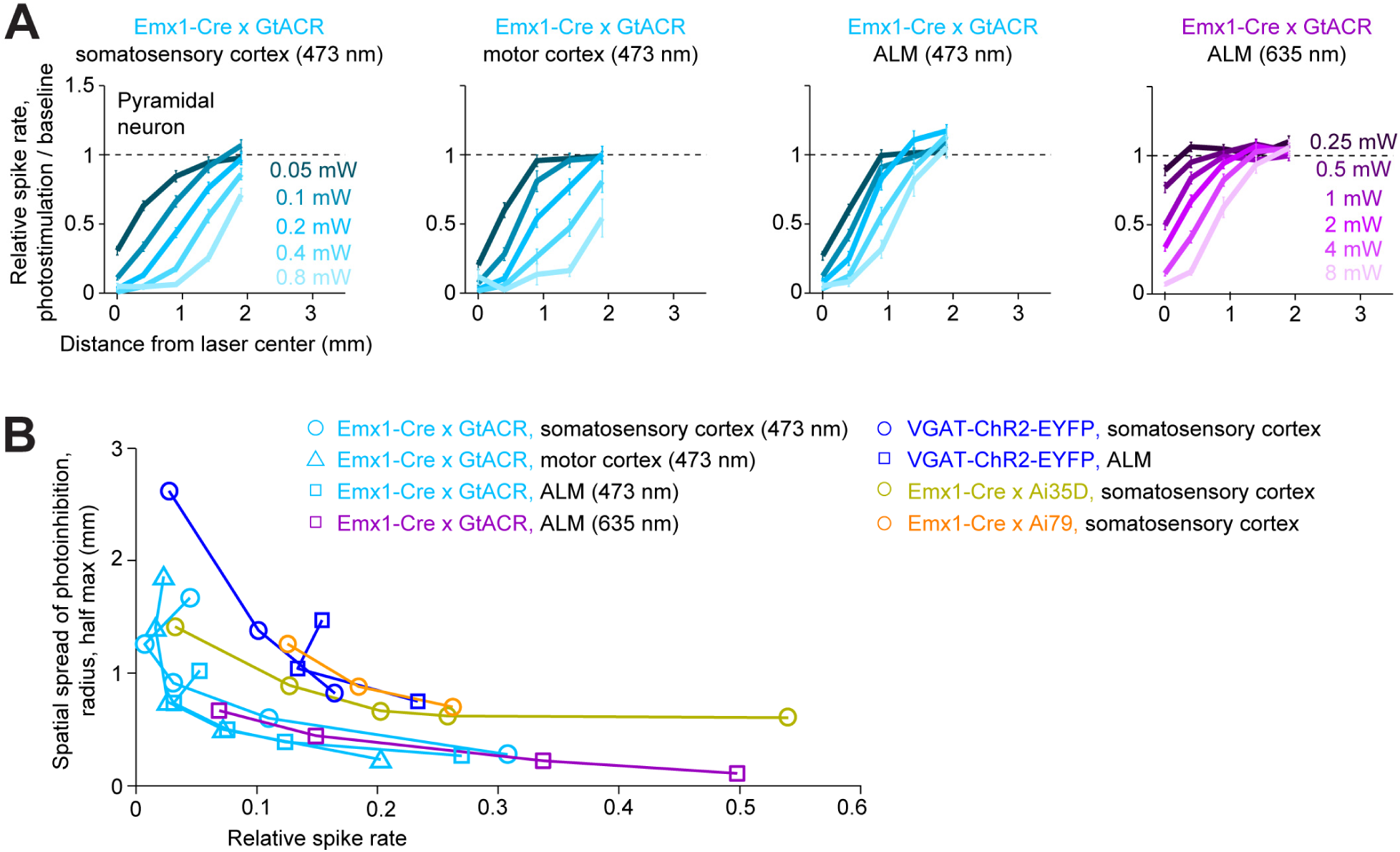
Spatial profile of direct photoinhibition in GtACR reporter mouse. (A) Normalized spike rate versus distance from the photostimulus center for various laser powers. Blue light (437 nm) photostimulation in somatosensory cortex, n=198 pyramidal neurons; blue light photostimulation in motor cortex, n=236; blue light photostimulation in ALM, n=335; red light (635 nm) photostimulation in ALM, n=236. Neurons were pooled across cortical depths. Mean ± sem, bootstrap across neurons. (B) Photoinhibition strength versus spatial spread. Normalized spike rate was averaged across all neurons near laser center (<0.4 mm, all cortical depths). Spatial spread is the distance at which photoinhibition strength was half of that at the laser center (“radius, half-max”). Each dot represents data from one photostimulation power. Lines connect all dots of one condition. VGAT-ChR2- EYFP, Emx1-Cre x Ai35D, Emx1-Cre x Camk2a-tTA x Ai79, data from Figure 5J replotted here for reference.

**Figure 7.**
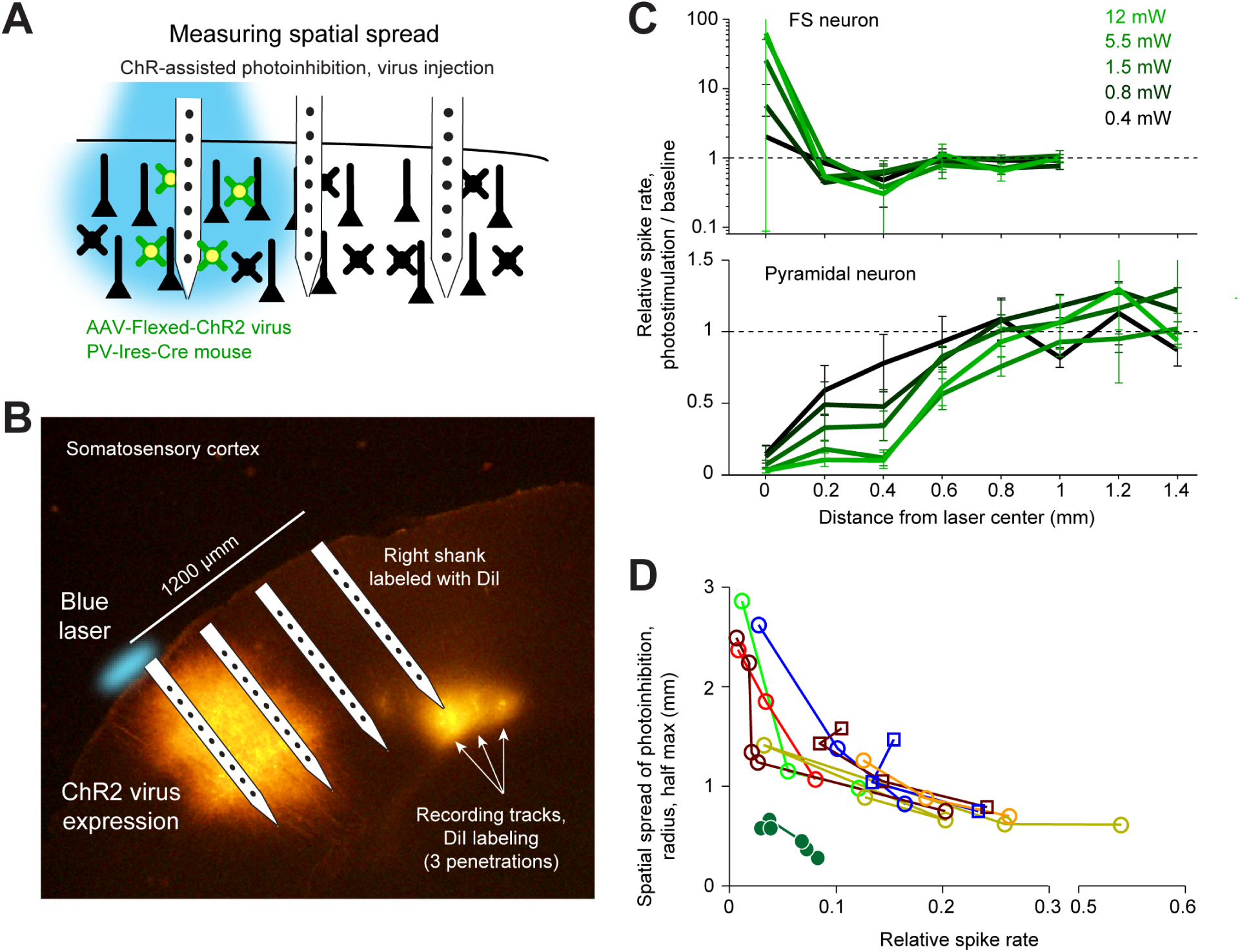
ChR-assisted photoinhibition using virus injection can achieve submillimeter spatial resolution. (A) Schematics, confined ChR2 expression in PV neurons and silicon probe recording at different distances from the expression site. (B) Silicone probe recording in barrel cortex during photostimulation. The right shank of the silicon probe was painted with DiI to label the recording tracks. Coronal section showing viral expression of ChR2-tdTomato, electrode and photostimulus locations. The photostimulus was aligned to the virus injection site. (C) Normalized spike rate versus distance from the photostimulus center for different laser powers. Top, FS neurons (n=14). Bottom, pyramidal neurons (n=78). Neurons were pooled across cortical depths. (D) Same as Figure 5J, but with virus injection data added.

We sought to limit the spatial spread of ChR-assisted photoinhibition by directly limiting the spatial profile of interneuron activation. We injected small volumes of Cre-dependent ChR2 virus in PV-IRES-Cre mice (Cardin et al., 2009; Lee et al., 2012; Lien and Scanziani, 2013; Pafundo et al., 2016). The virus injection localized the expression of ChR2 (diameter of expression, 500µm, Fig 7A). Silicon probe recordings (Fig 7B) confirmed that photostimulation excited FS neurons only at the infection site (Fig 7C). In the surrounding regions, FS neuron spike rates were even slightly suppressed (e.g. Fig 7C, at 400 µm away). Photoinhibition extended additional several hundred micrometers, with neurons suppressed even 800 µm away from the photostimulus center. When 90% of the spikes were removed at the infection site center, photoinhibition had a half-max radius of 0.5 mm (Fig 7D). Thus, limiting the spatial extend of interneuron excitation reduced photoinhibition spread to less than 1 mm (radius). These results show that optgogenetic inactivation methods have a fundamental resolution on the order of 1 mm (Fig 8).

**Figure 8.**
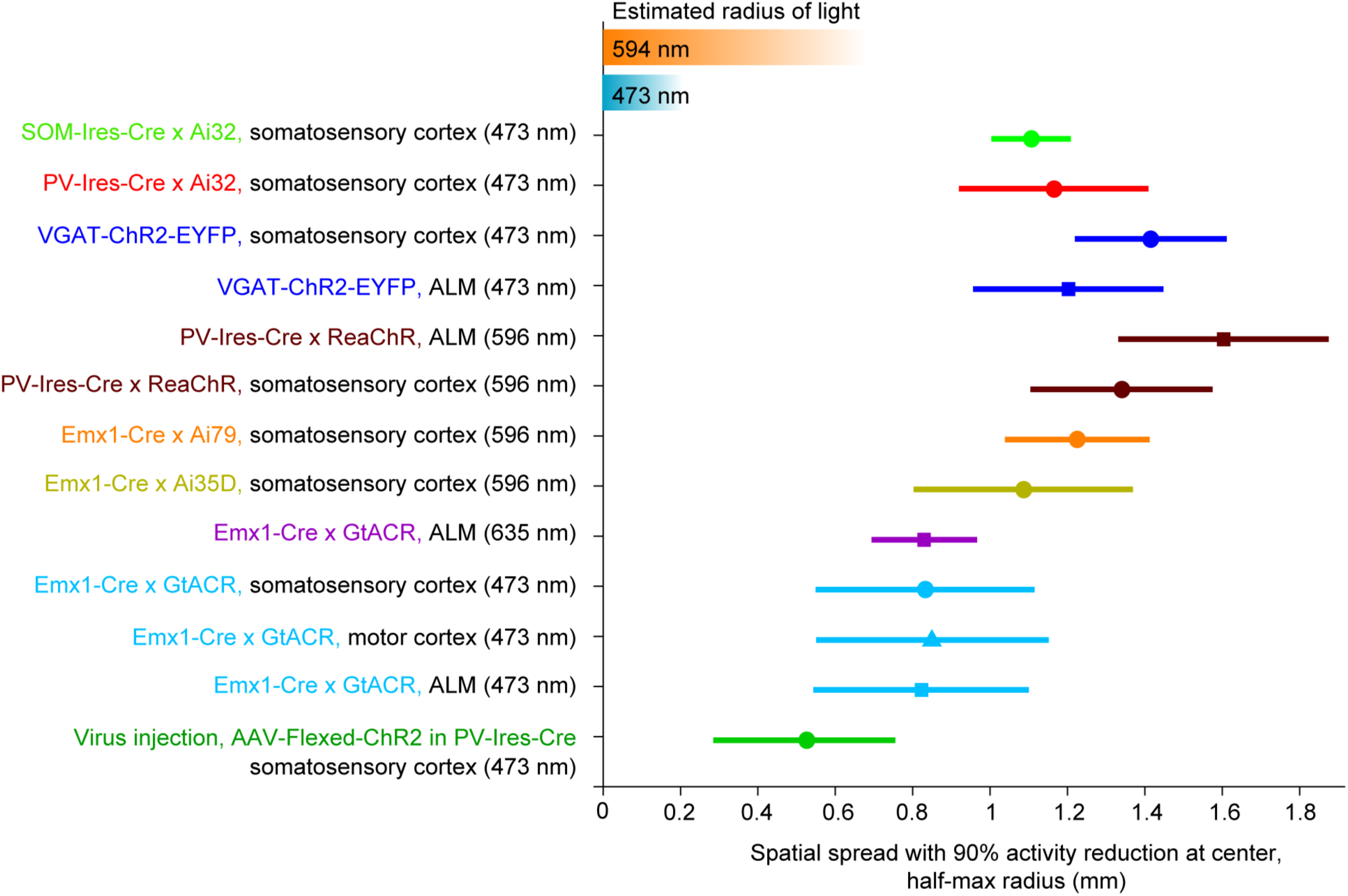
Summary of spatial resolution for all photoinhibition methods. Half-max radius of photoinhibition at 90% activity reduction at laser center. Data based on Figure 5J, 6B, and 7D. Mean ± sem, bootstrap across neurons.

### Strong coupling between cortical neurons and the paradoxical effect

What underlies the spread of photoinhibition? Cortical neurons are coupled with each other. Activity reduction in a small region in the vicinity of the photostimulus withdraws input to the surrounding regions, reducing activity in the surround. Consistent with this interpretation, in regions surrounding the photostimulation site, a concurrent decrease in activity was observed in both FS neurons and pyramidal neurons, even with ChR2-assisted photoinhibition (Fig 9, arrows). Activity decreased in proportion in FS and pyramidal neurons relative to their baseline activity.

**Figure 9.**
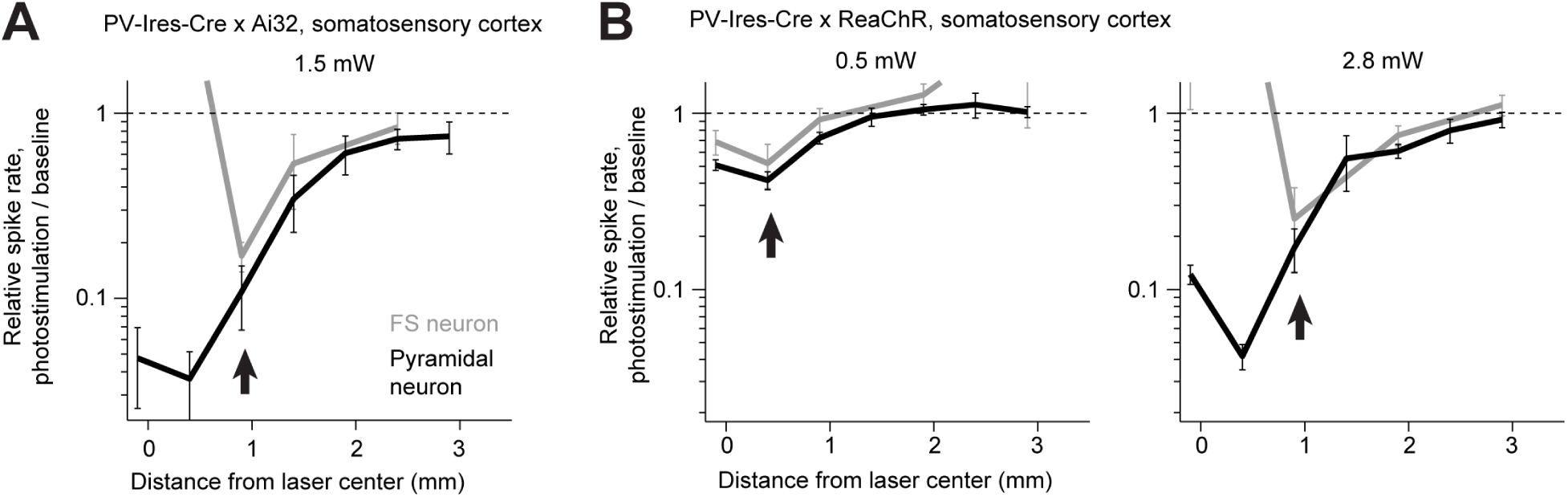
Proportional activity decrease in pyramidal and FS neurons in the photoinhibition surround during ChR2-assisted photoinhibition. (A) Normalized spike rate versus distance from the photostimulus center for PV-IRES-Cre x Ai32. Data from Figure 4B-D replotted with activity shown on a log scale. FS neurons (gray) and pyramidal neurons (black). The arrows point to regions in the surround where activity of FS neurons and pyramidal neurons decrease in proportion (paradoxical effect). (B) Same as (A) but for PV-IRES-Cre x ReaChR.

In standard models of cortical circuits, networks are stabilized by inhibition to prevent runaway excitation (Rubin et al., 2015; Tsodyks et al., 1997; van Vreeswijk and Sompolinsky, 1996) . These inhibition-stabilized networks (ISN) exhibit paradoxical effects. Selective excitation of interneurons decreases interneuron activity together with excitatory neurons (Litwin-Kumar et al., 2016; Pehlevan and Sompolinsky, 2014; Rubin et al., 2015; Tsodyks et al., 1997).

We tested this prediction in the somatosensory cortex. In mice expressing ReaChR in PV neurons, we used orange light to drive excitation in nearly all PV neurons across cortical layers (Fig 10A). Pyramidal neurons monotonically decreased their activity as a function of light intensity (Fig 10B). Interestingly, FS neurons also decreased their activity, despite being excited by ReaChR. As a function of light intensity, activity in FS neurons continued to decrease until most of the pyramidal neuron activity was silenced (Fig 10B, at 0.4 mW/mm^2^). At higher light intensities the FS neuron activity increased. This increase was partly driven by an increased photocurrent. At the same time, the network also transitioned to a regime with little coupling due to lack of excitatory activity. These data are consistent with the paradoxical effect predicted by cortical circuit models stabilized by inhibition (Tsodyks et al., 1997).

**Figure 10.**
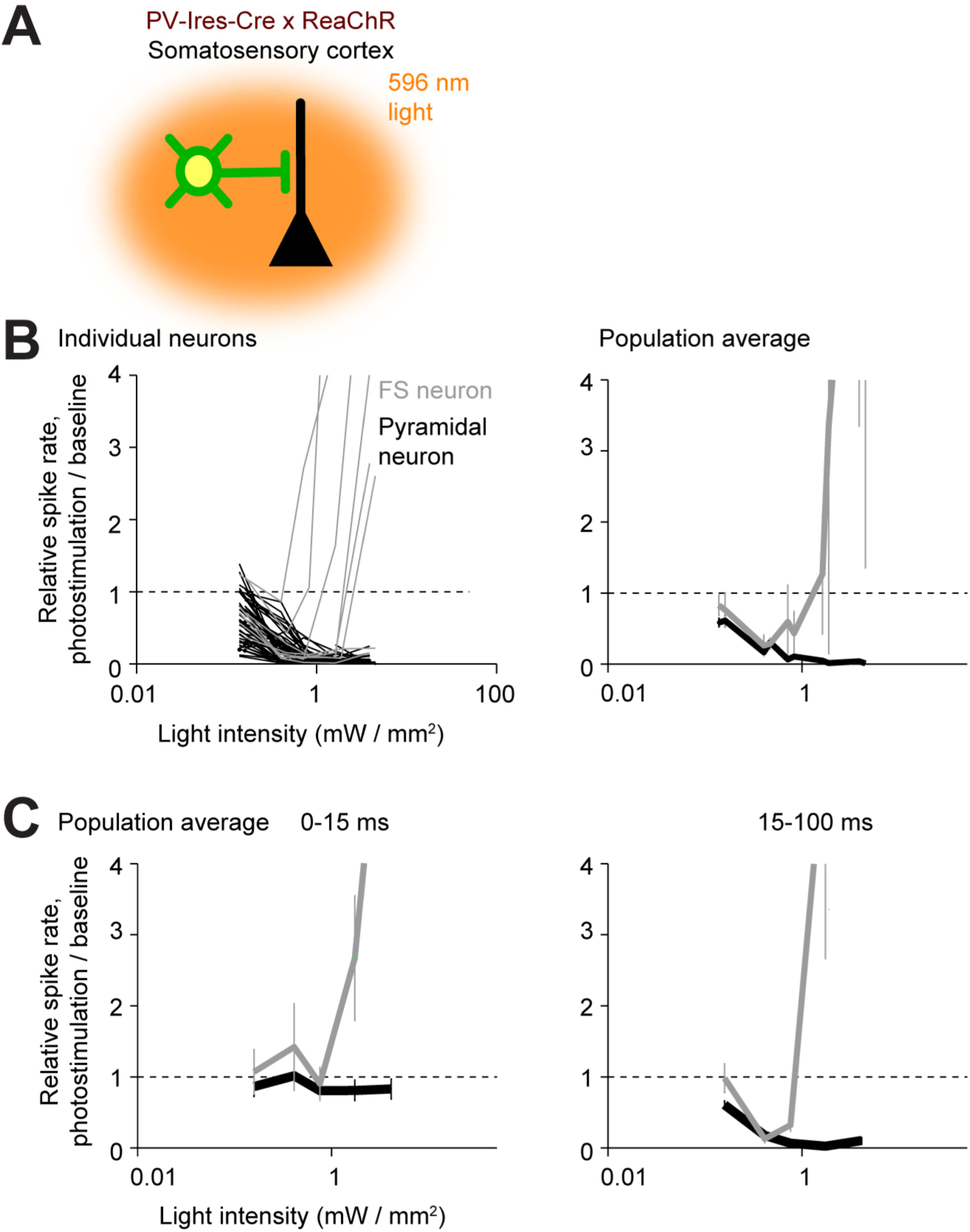
The paradoxical effect. (A) Photostimulating PV neurons using orange light. (B) Normalized spike rate as a function of light intensity (<0.4 mm from laser center, all cortical depths). FS neurons (red) and pyramidal neurons (blue). Top, individual neurons (lines). Bottom, mean ± s.e.m. across neurons, bootstrap. Laser power was divided by the illuminated area to obtain light intensity. FS neurons, n=10, pyramidal neurons n=82. (C) Same as (B) but for normalized spike rate at different epochs of photostimulation.

### Temporal profile of optogenetic inactivation

We examined the dynamics of photoinhibition. For ChR-assisted photoinhibition, the kinetics of ChR determined the dynamics of the interneurons (Fig 11A). For example, in mice expressing ChR2 (VGAT-ChR2-EYFP and PV-IRES-Cre x Ai32), FS neuron activity was time-locked to the photostimulus. In mice expressing ReaChR (PV-IRES-Cre x ReaChR), which has slower off kinetics compared to ChR2 (Lin et al., 2012), FS neuron activity was not time-locked to the photostimulus and was strongly attenuated over prolonged photostimulation. Despite the different interneuron dynamics, the photoinhibition of pyramidal neurons was similarly (Fig 11A). A subset of FS neurons were excited throughout the photostimulation (Fig 11B, 0.8-1 s). Other FS neurons were suppressed by photostimulation on average; however, a subset of these inhibited FS neurons were transiently excited by the photostimulus, followed by inhibition (Fig. 11B, compare 0-10 ms to 0.8-1 s), implying that these neurons were also expressing ChR. These data suggest that prolonged photostimulation (> 20 ms) triggered spikes in a subset of FS interneurons, which in turn reduced activity in other FS neurons and pyramidal neurons.

**Figure 11.**
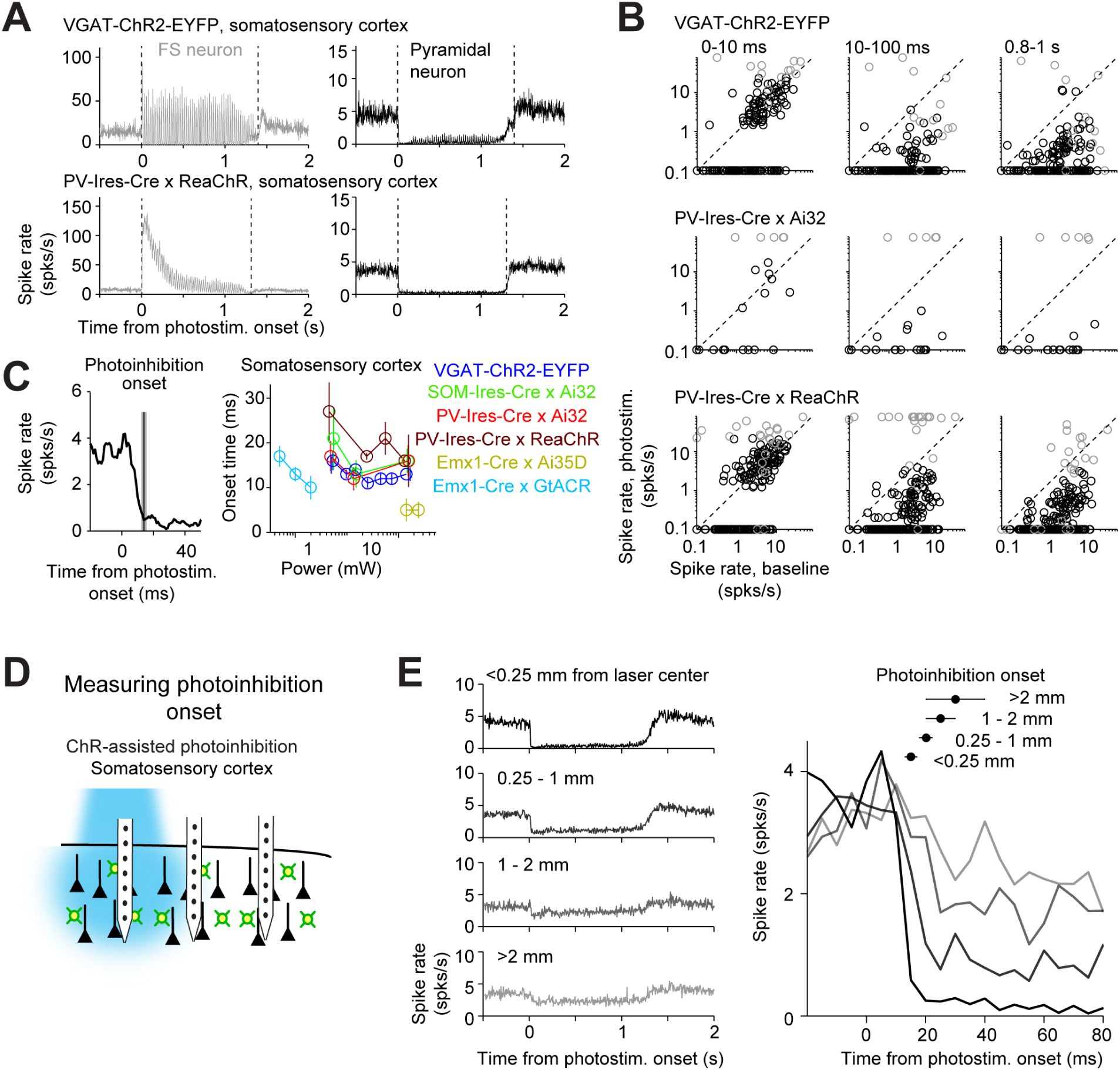
Time course of ChR2-assisted photoinhibition. (A) Mean peristimulus time histogram (PSTH, 1 ms bin) for FS neurons (gray) and pyramidal neurons (black). All neurons <0.4 mm from the laser center were pooled. Laser power, 14-15 mW. VGAT-ChR2-EYFP, FS neurons, n=12, pyramidal neurons n=152; PV-IRES-Cre x Ai32, FS neurons, n=5, pyramidal neurons n=16; PV-IRES-Cre x ReaChR, FS neurons, n=23, pyramidal neurons n=207. (B) Spike rate of FS neurons (gray) and pyramidal neurons (black) at different epochs of photostimulation. Dots correspond to individual neurons. Neurons with significant firing rate changes relative to baseline (*p* < 0.05, two-tailed *t*-test) are highlighted by circles. (C) Photoinhibition onset time. Left, PSTH of pyramidal neurons in VGAT-ChR2-EYFP mice and photoinhibition onset (mean ± sem). The photoinhibition onset is the time when spike rate reached 90% of the average spike rate reduction during the whole photostimulation period. Right, comparison of photoinhibition onset for different photoinhibition methods. Same color scheme as in Figure 1A. Each dot represents data from one photostimulation power. Lines connect all dots of one condition. (D) Schematic, measuring photoinhibition onset at different distances from the photostimulus. (E) Left, PSTH of pyramidal neurons at different distances from the photostimulus center. Right, spike rate after photoinhibition onset (t=0). Data from motor cortex and barrel cortex in VGAT-ChR2-EYFP, PV-IRES-Cre x Ai32, and PV-IRES-Cre x ReaChR mice are pooled (<0.25 mm, n=301; 0.25-1 mm, n=317; 1-2 mm, n=262; >2 mm, n=156). Laser power, 14-15 mW.

For ChR-assisted photoinhibition, the photoinhibition lagged FS neuron excitation by 3 ms (Materials and Methods, >1.5 mW). Photoinhibition was detectable 4.0 ± 1.7 ms after photostimulation onset (mean ±s.e.m., based on t-test of spike counts against baseline) and reached a maximum at 18.4 ± 1.6 ms (‘photoinhibition onset’, Fig 11C). ChR2-assisted photoinhibition has a slightly more rapid onset (14.6 ms) than ReachR-assisted photoinhibition (18.8 ms). Direct hyperpolarization of pyramidal neurons using Arch produced even more rapid photoinhibition onset (5 ms, Fig 11C).

Photoinhibition onset was progressively delayed at increasing distance from the laser center (Fig 11D-E). Photoinhibition at 2 mm away lagged the photoinhibition at the laser center by 10 ms (Fig 11E). This suggests a gradual spread of photoinhibition, consistent with the interpretation that coupling between cortical neurons mediated the photoinhibition spatial spread.

The temporal profile of photoinhibition offset was limited by rebound activity. Removal of photoinhibition elevated activity above baseline (‘rebound activity’), which decayed to baseline over hundreds of milliseconds (Fig. 12A). Rebound activity depended on both photostimulus duration and intensity (Fig 12B). For short photostimuli, rebound activity was moderate regardless of photostimulus intensity (Fig 12C, e.g. at 500 ms duration, rebound activity was 10% of baseline for 1.5-7mW). For long photostimulation durations, rebound activity was substantial even at lower laser power (Fig 12C, e.g. at 2 s duration, rebound activity was >20% of baseline at 1.5mW). At moderate photostimulation duration (1s), all photoinhibition methods produced some rebound activity regardless of photoinhibition strength (Fig 12D, Pearson’s correlation, r=-0.2, p=0.29). The rebound activity could be caused by recovery from synaptic depression in silenced excitatory neurons, synaptic depression in activated FS neurons, or perturbed intracellular ion concentrations accumulated over prolonged duration of photostimulation (Mahn et al., 2016; Wiegert et al., 2017).

**Figure 12.**
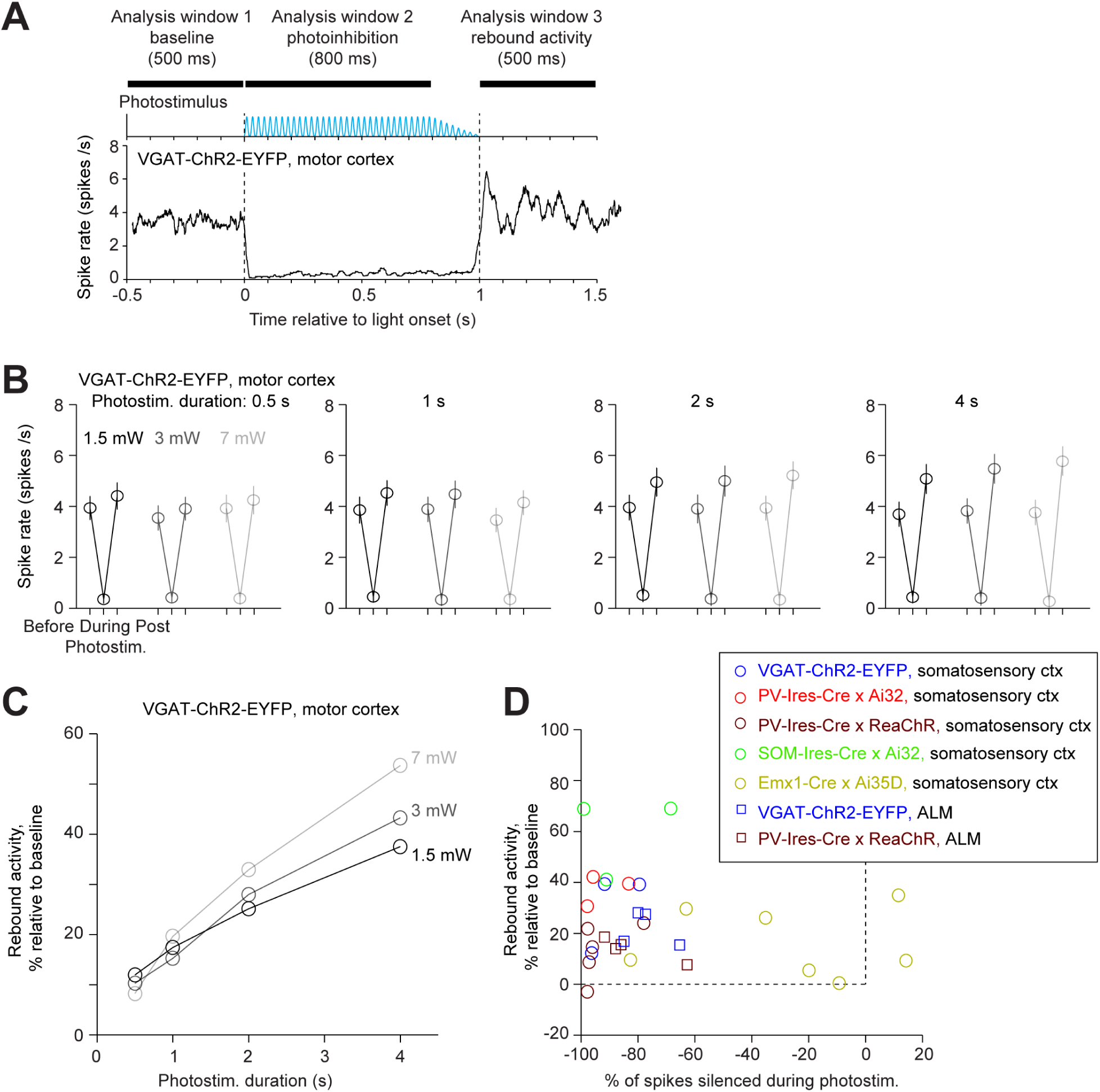
Rebound activity after photoinhibition. (A) PSTH of pyramidal neurons during photoinhibition in VGAT-ChR2-EYFP mice. All neurons <0.4 mm from the laser center (n=78). Photoinhibition was in motor cortex. Laser power, 7 mW. Spike rates were analyzed in 3 time windows before, during, and after photostimulation. (B) Spike rate of pyramidal neurons before, during, and after photostimulation for different laser powers and photostimulation durations. (C) Rebound activity as a function of laser powers and photostimulation durations. Relative activity is the post-photostimulation spike rate increase from the baseline, normalized to the baseline spike rate. (D) Rebound activity as a function of photoinhibition strength. Percent of spikes silenced was relative to the baseline spike rate. Data from all mouse lines. VGAT-ChR2-EYFP, somatosensory cortex, n=111; PV-IRES-Cre x Ai32, somatosensory cortex, n=16; PV-IRES-Cre x ReaChR, somatosensory cortex, n=82; SOM-IRES-Cre x Ai32, somatosensory cortex, n=65; Emx1-Cre x Ai35D, somatosensory cortex, n=26; VGAT-ChR2-EYFP, ALM, n=96; PV-IRES-Cre x ReaChR, ALM, n=129.

In summary, photoinhibition onset was rapid (10 ms), but the offset was limited by rebound activity. Attenuating the photostimulus gradually (100ms) or reducing the photostimulus durations can both reduce rebound activity (Fig 12). These factors impose constraints on the temporal resolution of photoinhibition.

## Discussion

We examined optogenetic methods to locally silence neural activity in the mouse neocortex. ChR-assisted photoinhibition was effective at lower light intensities in suppressing local pyramidal neurons than direct photoinhibition using the light-gated ion pumps Arch and Jaws. A soma localized Cl-channel, GtACR1 (Govorunova et al., 2015; Mahn et al., 2018), produced the strongest photoinhibition. We generated a reporter mouse that expresses the soma localized GtACR1 (Mahn et al., 2018) under the control of Cre-recombinases. This transgenic mouse will be a useful tool to silence activity in genetically defined neuron populations.

We characterized the spatial profile of different photoinhibition methods. Photoinhibition spread over 1 mm or more, far beyond the spatial spread of light (0.25 – 0.5 mm). The spatial extent of photoinhibition was similar in somatosensory and motor cortex despite differences in microcircuits and connections with thalamus, and similar across photoinhibition methods (Fig 5-8). The spatial profiles of photoinhibition, laterally and along the beam direction, were also similar across different wavelengths (473 vs. 593 nm), despite large differences in the spatial profiles of the photostimuli (Fig 3).

Local infection with Cre-dependent, ChR2-expressing virus in PV-IRES-Cre mice produced the smallest lateral spread in photoinhibition (∼0.5 mm in radius). Local expression of ChR limited the excitation of FS interneurons to a small region (∼200 um in radius, Fig 7C). Additional lateral spread of photoinhibition was produced by the cortical circuit. Given the limited penetration of light in tissue, the direct effects of photostimulation on neurons were most pronounced in L2/3. Silencing activity in a focal region in the upper layers caused a withdrawal of excitatory input to deeper layers (Hooks et al., 2011; Kiritani et al., 2012) and to surrounding cortical regions (Lefort et al., 2009; Ozeki et al., 2009). For example, in regions surrounding the viral injection site, photostimulation caused a decrease in activity of both FS neurons and pyramidal neurons. Future studies directly recording inhibition onto neurons in the surround could further clarify the underlying network effect.

In both sensory and motor cortex, cortical neurons cannot maintain activity without thalamic drive (Guo et al., 2017; Reinhold et al., 2015). Conversely, silencing activity in cortex reduces activity in the thalamus (Guo et al., 2017). Therefore, silencing a region in cortex likely decreased activity in the parts of thalamus targeted by the silenced region. Since corticothalamic projections arise from the deep layers, and the direct effects of photositmulation occurred mostly in superficial layers (with blue light), our data imply that intracortical connections are critical to maintain cortical activity, in addition to thalamocortical input.

Our data provides additional evidence for strong coupling between cortical neurons. Weak excitation of interneurons induced the “paradoxical effect”, where activity decreased in both interneurons and excitatory neurons (Fig 10). Excitation of GABAergic neurons reduces activity in nearby excitatory neurons, and thus reduced excitation to GABAergic neurons. Paradoxical effects were also observed elsewhere in our experiments. For example, in Fig 5B-D at laser center, weak photostimulation (0.5 mW) of GABAergic neurons induced little increase or even a decrease in FS neurons activity, yet, pyramidal neuron activity was silenced. These observations are consistent with predictions of inhibition-stabilized networks, and they suggest that cortical neurons operate in a regime with strong coupling (Lefort et al., 2009; Ozeki et al., 2009; Pehlevan and Sompolinsky, 2014; Rubin et al., 2015; Tsodyks et al., 1997).

Previous studies manipulated cortical interneurons but did not find the paradoxical effect predicted by inhibition-stabilized network models (Atallah et al., 2012; Gutnisky et al., 2017; Yu et al., 2016). One key difference is that we weakly photostimulated nearly all PV neurons in superficial layers of transgenic mice. Previous studies used viral expression strategies that manipulated only subsets of PV neurons. Theoretical analysis and simulations suggest that the paradoxical effect is only induced when a large proportion of interneurons is excited (Gutnisky et al., 2017; Sadeh et al., 2017). Another key difference is photostimulation power. Cortical networks escape the paradoxical effect regime when most of the pyramidal neurons are silenced (Fig 10B), which effectively removes coupling between excitatory neurons. Consistent with this prediction, the paradoxical effect was only be observed under weak photostimulation conditions that do not silence a majority of pyramidal neurons (Fig. 10).

Our data provide guidance on the design of *in vivo* optogenetic experiments. Photoinhibition has been used extensively in loss-of-function studies to localize functions to specific brain regions. In the neocortex, the 1 mm length scale of photoinhibition poses a fundamental limit on the spatial resolution of loss-of-function manipulations. Interpretation of photoinhibition effects must take this spatial spread function into account. For example, silencing a brain region nearby to an involved region could produce false-positive behavioral effects. To get around this, one could systematically map the photoinhibition effects around a region of interest, then deconvolve the known spatial spread function to recover the underlying region involved in behavior (Li et al., 2016).

We note that the spatial resolution of photoinhibition could differ across brain regions. In many subcortical regions GABAergic neurons make long-range projections, reducing the spatial resolution of ChR2-assisted photoinhibition. On the other hand, recurrent and lateral excitation is less pronounced in many subcortical regions compared to neocortex, promising better spatial resolution when direct photoinhibition is used for photoinhibition.

Photoinhibition has been used to probe the involvement of brain regions during specific behavioral epochs (Guo et al., 2014b; Hanks et al., 2015; Li et al., 2015; Sachidhanandam et al., 2013). Our results suggest that the temporal resolution of photoinhibition was limited by rebound activity (Fig 12), which could potentially produce confounding behavioral consequences. Rebound activity could be partially alleviated by changes in photostimulus parameters (Fig 12C) (Wiegert et al., 2017).

In many experiments, cell-type specific manipulations produced multi-phasic network responses both on local and downstream circuits over hundreds of milliseconds (Guo et al., 2017). These results highlight that optogenetic circuit manipulations provide the most insights into functions of specific neural dynamics when the primary effects of the perturbation and, equally importantly, the effects on downstream brain areas are taken into account, ideally measured simultaneously in behaving animals (Gao et al., 2018; Guo et al., 2017; Inagaki et al., 2019; Li et al., 2016).

## Acknowledgements.

We thank Mathias Mahn and Ofer Yizhar for sharing the GtACR1-ts-FRed-Kv2.1 construct. We thank Mingshan Xue for comments on the manuscript, David Golomb for critical discussions, Susan Michael and Amy Hu for histology, Tim Harris, Brian Barbarits, Anthony Leonardo for help with silicon probe recordings. This work was supported by Howard Hughes Medical Institute (K.S.), Helen Hay Whitney Foundation fellowship (N.L., H.I.), Sir Henry Wellcome Postdoctoral Fellowship (S.C.), the Robert and Janice McNair Foundation (N.L.), Whitehall Foundation (N.L.), Alfred P. Sloan Foundation (N.L.), Searle Scholars Program (N.L.), NIH NS104781 (N.L.), the Pew Charitable Trusts (N.L.), and Simons Collaboration on the Global Brain (#543005, K.S., N.L.).

## Author Contributions

NL, KS conceived the project. NL, SC, ZVG, HC, YH, HI, and CD performed the experiments. NL, SC, ZVG, HC, YH, HI, and KS analyzed the data. CG and HI generated the GtACR reporter mouse. SC and HI performed characterizations of the GtACR1 reporter mouse. NL and KS wrote the paper with comments from other authors.

## Materials and Methods

### Animals

This study is based on data from 39 mice (age > P60, both male and female mice). 29 transgenic mice were used to characterize photoinhibition, including 14 VGAT-ChR2-EYFP mice, 2 PV-IRES-Cre x Ai32 mice, 3 SOM-IRES-Cre x Ai32 mice, 5 PV-IRES-Cre x R26-CAG-LSL-ReaChR-mCitrine mice, 1 Emx1-Cre x Ai35D mouse, 2 Emx1-Cre x Camk2a-tTa x Ai79 mice, and 2 Emx1-Cre x R26-CAGLNL-GtACR1-ts-FRed-Kv2.1 mice. 5 PV-IRES-Cre mice were used to drive photoinhibition using Cre-dependent ChR2 virus injections. 2 Tlx_PL56-Cre x Ai32 mice and 1 Rbp4-Cre x R26-CAG-LSL-ArchT-GFP mouse were used to characterize cell-type specific manipulations. 1 Emx1-Cre x Rosa26-LSL-H2B-mCherry mouse and 1 Emx1-Cre x Rosa-CAG-LSL-H2B-GFP mouse (gift from Josh Huang, Cold Spring Harbor Laboratory) were used for photobleaching experiments. All procedures were in accordance with protocols approved by the Janelia Farm Institutional Animal Care and Use Committee.

### Generation of transgenic GtACR mice

Targeting vector, Rosa26-CAG-LNL-GtACR1-ts-FusionRed-Kv2.1C-WPRE-polyA, was derived from AAV.CamKIIa.GtACR1-ts-FRed -Kv2.1.WPRE (Mahn et al., 2018) and a standard Rosa26 backbone (Dymecki and Kim, 2007; Madisen et al., 2012; Soriano, 1999). The targeting vector DNA was electroporated into ES cells and the chimeric mice were generated using successfully targeted ES cell clones. GtACR expression was evaluated both by native fluorescence and functional assays. Rosa26-CAG-LNL-GtACR1-ts-FRed-Kv2.1 mice were submitted to The Jackson Laboratory (stock #033089).

### Surgery

Mice were prepared for photostimulation and electrophysiology with a clear-skull cap (Guo et al., 2014b) and a headpost (Guo et al., 2014a). The scalp and periosteum over the dorsal surface of the skull were removed. A layer of cyanoacrylate adhesive (Krazy glue, Elmer’s Products Inc) was directly applied to the intact skull. A custom made headbar was placed on the skull (approximately over visual cortex) and cemented in place with clear dental acrylic (Lang Dental Jet Repair Acrylic; Part# 1223-clear). A thin layer of clear dental acrylic was applied over the cyanoacrylate adhesive covering the entire exposed skull, followed by a thin layer of clear nail polish (Electron Microscopy Sciences, Part# 72180).

In some PV-IRES-Cre mice ChR2 was introduced by injecting 50 nL of AAV2/5-hSyn1-FLEX-hChR2-tdTomato (Addgene plasmid 41015, Janelia Molecular Biology Shared Resource) (O’Connor et al., 2013) into the barrel cortex (bregma posterior 1 mm, lateral 3.5 mm) at two depths (400 and 800 µm), followed by implantation of the headbar. The injection was made through the thinned skull using a custom, piston-based, volumetric injection system. Glass pipettes (Drummond) were pulled and bevelled to a sharp tip (outer diameter of 30 µm) (Petreanu et al., 2009). Pipettes were back-filled with mineral oil and front-loaded with viral suspension immediately before injection.

For silicon probe recordings, a small craniotomy was made over the recording site in mice already implanted with the clear-skull cap and headpost (see *Electrophysiology*). The dental acrylic and bone were thinned using a dental drill. The remaining thinned bone was carefully removed using a bent forceps. A separate, smaller craniotomy (diameter, approximately 600 µm) was made through the headpost bar for ground wIRES.

### Photostimulation

Light from a 473 nm laser (DHOM-M-473-200, UltraLaser) or a 594 nm laser (Cobolt Inc., Colbolt Mambo 100) or a 635 nm laser (MRL-III-635L-100mW, Changchun New Industries Optoelectronics Technology) was controlled by an acousto-optic modulator (AOM; MTS110-A3-VIS, Quanta Tech; extinction ratio 1:2000; 1µs rise time) and a shutter (Vincent Associates), coupled to a 2D scanning galvo system (GVSM002, Thorlabs), then focused onto the brain surface (Guo et al., 2014b). The laser at the brain surface had a Gaussian profile with a beam diameter of 400 µm at 4σ. We tested photoinhibition in barrel cortex (bregma posterior 0.5 mm, 3.5 mm lateral), anterior lateral motor cortex (ALM, bregma anterior 2.5mm, 1.5 mm lateral) and primary motor cortex (bregma anterior 0.5mm, 1.5 mm lateral). Photoinhibition was similar across different regions.

To prevent the mice from detecting the photostimulus, a ‘masking flash’ (40 x 1 ms pulses at 10 Hz) was delivered using a LED driver (Mightex, SLA-1200-2) and 470 nm LEDs (Luxeon Star). The masking flash began before the photostimulus started and continued through the end of the epoch in which photostimulation could occur.

The standard photostimulus had a near sinusoidal temporal profile (40 Hz) with a linear attenuation in intensity over the last 100-200 ms (duration: 1.3 s including the ramp). This temporal profile was chosen to minimize rebound activity based on pilot experiments. In some cases, we also used a constant photostimulus without ramp (Fig 1). In the experiments presented in Figure 12 we also tested other photostimulus durations (0.5, 1, 2, and 4 s including the ramp). The photostimuli were delivered at approximately 7 s intervals. The power (0.53, 1.0, 1.5, 2.5, 4.5, 7.3, 14 mW) and locations of photostimulation (0.5, 0.75, 1.0, 2.0, 3.0 mm from the recording sites) were chosen randomly. Because we used a time-varying photostimuli the power values reported here reflect the time-averaged power.

### Photobleaching

We used the rate of photobleaching *in vivo* to measure the spatial profile of light intensity at two different wavelengths (473 nm and 594 nm, Fig 3). In transgenic mice expressing GFP (absorption at 473 nm) or mCherry (absorption at 594 nm) in the nuclei of cortical pyramidal neurons, photobleaching was induced by prolonged (10 min) illumination at different laser powers through the clear-skull cap. Nuclear fluorescence was imaged in fixed tissue sections using a confocal microscope (Zeiss LSM 510). Multiple sections were imaged around each photobleaching site.

GFP or mCherry fluorescence was measured in regions of interest (ROIs) around individual nuclei (Fig 3). For each ROI, we calculated a ΔF by substracting a baseline ROI intensity, F_0_, from the mean fluorescence. The F_0_ was computed by averaging the intensity of all ROIs > 700 µm away from the laser center. Near the photostimulus center, photobleaching was measured as the average ΔF/ F_0_ of all ROIs within a small region near the photostimulus center (within 100 µm for GFP, 400 µm for mCherry). Photobleaching (ΔF/ F_0_) monotonically increased with laser power. We used the empirical relationship between ΔF/ F_0_ and laser power to convert GFP or mCherry intensities into power to obtain the photostimulus spatial profile (Fig 3D, G).

### Electrophysiology

All recording were carried out while mice were awake but non-behaving. Extracellular spiking activity was recorded using silicon probes. We used NeuroNexus probes with 4 shanks (at 200 or 400 µm spacing) and recording sites spaced 100 or 200 µm apart (8 sites per shank, P/N A4x8-5mm-100-200-177, A4x8-5mm-100-200-413, A4x8-5mm-200-200-177, A4x8-5mm-200-200-413, and, A4x8-5mm-100-400-177). Silicon probes were connected to a headstage (Intan Technology) that multiplexed the 32-channel voltage recording into 2 analog signals (fabricated at Janelia Farm Research Campus, Brian Barbarits, Tim Harris). The multiplexed analog signals were recorded on a PCI6133 board at 312.5 kHz (National instrument) and digitized at 14 bit. The signals were demultiplexed into the 32 voltage traces at the sampling frequency of 19531.25Hz and stored for offline analyses in a custom software spikeGL (C. Culianu, Anthony Leonardo, Janelia Farm Research Campus). The headstage was mounted on a motorized micromanipulator (MP-285, Sutter Instrument).

A 1 mm diameter craniotomy over the recording site was made prior to the recording sessions. In PV-IRES-Cre mice prepared for ChR2 virus mediated photoinhibition, a larger craniotomy (2 mm x 1 mm) was made around the injection site. The position of the craniotomy was guided by stereotactic coordinates for recordings in ALM (bregma anterior 2.5mm, 1.5 mm lateral), motor cortex (bregma anterior 0.5mm, 1.5 mm lateral), or barrel cortex (bregma posterior 0.5 mm, 3.5 mm lateral).

Prior to each recording session, the tips of the silicon probe were brushed with DiI in ethanol solution and allowed to dry. The surface of the craniotomy was kept moist with saline. The silicon probe was positioned on the surface of the cortex and advanced manually into the brain at ∼ 3 µm/s, normal to the pial surface. The depth of the electrode tip ranged from 749 to 1000 µm below the pial surface. The electrode depth was inferred from manipulator depth and verified with histology. To minimize pulsation of the brain, a drop of silicone gel (3-4680, Dow Corning, Midland, MI) was applied over the craniotomy after the electrode was in position. The tissue was allowed to settle for several minutes before the recording was started.

### Histology

After the conclusion of recording experiments, mice were perfused transcardially with PBS followed by 4% PFA / 0.1 M PB. The brains were fixed overnight and sectioned. Coronal sections (100 µm) were cut and images of DiI labeled recording tracks were acquired on a macroscope (Olympus MVX10). Electrode tracks were compared to the manipulator depth readings (Guo et al., 2014b).

### Data analysis

The extracellular recording traces were band-pass filtered (300-6 kHz). Events that exceeded an amplitude threshold (4 standard deviations of the background) were subjected to manual spike sorting to extract single-units (Guo et al., 2014b).

Our final data set comprised of 2571 single units (barrel cortex, 1158; ALM, 1038; M1, 375). For each unit, its spike width was computed as the trough to peak interval in the mean spike waveform (Fig 1B). We defined units with spike width <0.35 ms as FS neurons (354/2571) and units with spike width >0.45 ms as putative pyramidal neurons (2151/2571). Units with intermediate values (0.35 - 0.45 ms, 36/2571) were excluded from our analyses.

To quantify the strength of inactivation, we derived a “normalized spike rate”. For each neuron, we computed its spike rate during the photostimulus and its baseline spike rate (500 ms time window before photostimulus onset). The spike rates with photostimulation were averaged across the population (without normalization) and normalized by dividing the averaged baseline spike rate. The “normalized spike rate” reports the spike rate left during photostimulation.

To quantify the spatial spread of inactivation we computed normalized spike rate at different distances from the photostimulus, and the distance at which inactivation is half of max, that is its strength at the photostimulus center (Fig 5J, “radius, half-max”).

To quantify the time course of inactivation, we computed the average population Peristimulus Time Histogram (PSTH) aligned to photostimulus onset (Fig. 11). To quantify the onset time of inactivation, we estimated the time when the pyramidal neuron PSTH reached its minimum post stimulus onset (18.4 ± 1.6 ms, mean ± s.e.m.). To quantify the earliest time inactivation occurred (3.95 ± 1.7 ms), we estimated the first time bin (10 ms bin) in which the spike rate differed significantly (p < 0.05 two-tailed t-test) from baseline. To quantify photoinhibition lag from excitation of the interneurons, we computed the earliest time bin in which FS neuron PSTH significantly deviated from baseline spike rate (1.05 ± 0.2 ms), and subtracted this onset time from the photoinhibition onset time (lag: 2.90 ms). Bootstrap was performed over neurons to obtain standard error of the mean.

To quantify rebound activity, for each neuron, we computed spike rate in a 500 ms window after photostimulus offset, and normalized this spike rate to its baseline spike rate (500 ms time window before photostimulus onset). The rebound activity was quantified as the percentage of activity increase from baseline. The rebound activity was averaged across the recorded population (Fig 12C-D).

